# Climate-Change Driven Decline of an Insect Pathogen Increases the Risk of Defoliation by a Forest Pest Insect

**DOI:** 10.1101/2023.11.01.564627

**Authors:** Jiawei Liu, Colin Kyle, Jiali Wang, Rao Kotamarthi, William Koval, Greg Dwyer

## Abstract

The effects of climate change on forest-defoliating insects are poorly understood, a problem that is especially urgent in the case of the spongy moth (formerly “the gypsy moth”, *Lymantria dispar*). For decades following its introduction in 1869, the spongy moth severely defoliated North American forests, but the introduction of the pathogen *Entomophaga maimaiga* in 1989 drastically lowered defoliation levels. *E. maimaiga*, however, needs cool, moist conditions, whereas climate change is bringing hot, dry conditions to the range of the spongy moth. Here we use an empirically validated mathematical model to project that climate change will sharply reduce *E. maimaiga* infection rates, greatly increasing spongy moth defoliation. Recent data show that defoliation has strongly rebounded, supporting our projections. Our work shows that the effects of climate change on insect pathogens can have dire consequences for forests, and demonstrates the importance of understanding how climate change can alter species interactions.

---

Populations of many insect species have been decimated by climate change, leading to significant losses of insect diversity [1]. In some species, however, climate change has led to increased population sizes [2]; populations of tree-girdling bark beetles, for example, have increased because of the lengthened growing seasons that result from increased temperatures [3], leading to sharp increases in tree mortality [4]. Overall trends of negative effects of climate change on insects in general thus do not preclude positive effects of climate change on forest insect pests in particular.

Forest-defoliating insects also cause severe damage to forests [5], but the extent to which forest defoliators are affected by climate change is not well understood [6]. Recent climate anomalies temporarily reduced outbreak severity in the defoliating larch budmoth, *Zeiraphera diniana* [7], but projecting the effects of climate change on defoliators is difficult because many defoliators, including the larch budmoth, are held in check by specialist pathogens [8] and parasitoids [9]. Projecting how climate change will affect forest defoliators, and thus forests, therefore requires that we project the effects of climate change on interactions between forest trees, forest defoliators and the pathogens and parasitoids of forest defoliators.

Here we project the effects of climate change on the interaction between North American hardwood trees, the invasive spongy moth (*Lymantria dispar*, formerly “the gypsy moth”), and the spongy moth’s fatal, specialist, fungal pathogen *Entomophaga maimaiga*. For decades following the insect’s introduction into North America in 1869, the annual costs of spongy moth defoliation were in the millions of US dollars [10], but the introduction of *E. maimaiga* in 1989 greatly reduced defoliation [11]. Like many fungal pathogens [12], however, *E. maimaiga* spreads more rapidly when conditions are cool and moist [13], and so the hot, relatively dry conditions that climate change is bringing to the insect’s range in eastern North America [14] may lead to the resurgence of the spongy moth.

Moreover, a range-change model has projected that climate change will allow the spongy moth to invade areas that are beyond the northern edge of its range [15], leading to defoliation in new regions. The range-change model, however, considers only the effects of climate change on spongy moth growth rates, whereas defoliation levels depend as well on spongy moth densities [16], which in turn depend on *E. maimaiga* densities. Projecting defoliation rates therefore required that we construct a model that could project the densities of both the spongy moth and *E. maimaiga*.

## Model Construction

Ecological models of species interactions, like the spongy moth-*E. maimaiga* interac-tion, are expressly designed to project the densities of interacting species [17], but the use of species interaction models to project the ecological effects of climate change has been hindered by two obstacles. First, empirical studies usually consider only how climate change will affect individual species [18]; even when species interactions are taken into account, the high data requirements of the models means that existing data are often insufficient to allow model parameterization. If sufficient data are available, a second obstacle is that the allowance for multiple species leads to nonlinear terms in the models, and so parameterizing the models requires the implementation of non-linear fitting algorithms [19]. In previous work, however, three of us and colleagues collected a data set on the effects of weather on *E. maimaiga* spread rates in spongy moth populations that was sufficient for model parameterization, and we combined our data with a nonlinear fitting algorithm to estimate the parameters of a weather-driven model of the spongy moth-*E. maimaiga* interaction[20].

The data to which we fit our model consist of estimates of *E. maimaiga* infection rates in larval spongy moth populations (only larvae are susceptible to the pathogen) in the years 2010-2012 at a range of spongy moth densities along a 300 km-long north/south transect in Michigan, USA. Weather conditions in our study were similar to weather conditions across the range of the spongy moth in eastern North America during the same period (Supplemental Information), and so here we use our model to make projections across the entire North American range of the insect. To focus on forests known to be suitable for the spongy moth, we make projections only in areas that have previously experienced spongy moth defoliation (Supplemental Information; Canadian defoliation data are only publicly available for Ontario).

Efforts to project the effects of climate change on species interactions [21, 22] have often parameterized models using experiments in which organisms are grown at constant temperatures in the laboratory. Because we instead parameterized our model using field data, we allowed for multiple weather variables and realistic weather conditions. To identify underlying mechanisms, we fit our models to a combination of experimental and observational data [20], and we used statistical model selection [23] to choose between fifteen models that made different assumptions about the *E. maimaiga*-spongy moth interaction. Because it is of course possible that our model would not provide as good a fit to other data sets, in the Supplemental Information we compare the model’s projections to literature data. Although previous studies did not collect enough data for us to carry out the kind of statistically robust test that we used in our previous work, the data are nevertheless sufficient to carry out at least a qualitative test of our model, a test that the model easily passes.

The best model includes effects of temperature, relative humidity, rainfall, and the densities of the spongy moth and *E. maimaiga*, whereas models that did not include spongy moth and *E. maimaiga* densities, or that included only two, one or no weather variables, provided substantially worse explanations for the data [20]. The dynamics of our model are thus density-dependent, so that high spongy moth densities lead to high *E. maimaiga* infection rates, as occurs in nature [20]. If weather conditions in nature or in the model are sufficiently cool and moist, however, *E. maimaiga* transmission can be high even if the spongy moth population density is low. Since the introduction of *E. maimaiga*, weather conditions in eastern North America have indeed been cool enough and moist enough that *E. maimaiga* has often been able to decimate low and intermediate density spongy moth populations [13], preventing severe defoliation. The high temperatures and low relative humidity that climate change is bringing to eastern North America [14] nevertheless mean that climate change may allow the spongy moth to again become a major pest of hardwood forests.

## Model Projections

To project how climate change will alter weather conditions at ecologically relevant scales, we used a technique known as “dynamic down-scaling”, in which continental-scale projections from standard climate-change models are converted to the forest scales at which *E. maimaiga* epizootics occur [14, 24] (an epizootic is an epidemic in an animal population). Our dynamic downscaling simulations project that temperatures will increase and relative humidity will decrease at the vast majority of locations in eastern North America, but also project that rainfall will increase in almost as many locations as it will decrease (fig. 1). Because increased rainfall has positive effects on *E. maimaiga*, while increased temperatures and decreased relative humidity have negative effects on *E. maimaiga* [20], the climate-change model alone cannot be used to project whether climate change will have positive or negative effects on *E. maimaiga*. To make such projections, we therefore combined our ecological model with our climate-change model to form what we call an “eco-climate” model.

**Fig. 1.**
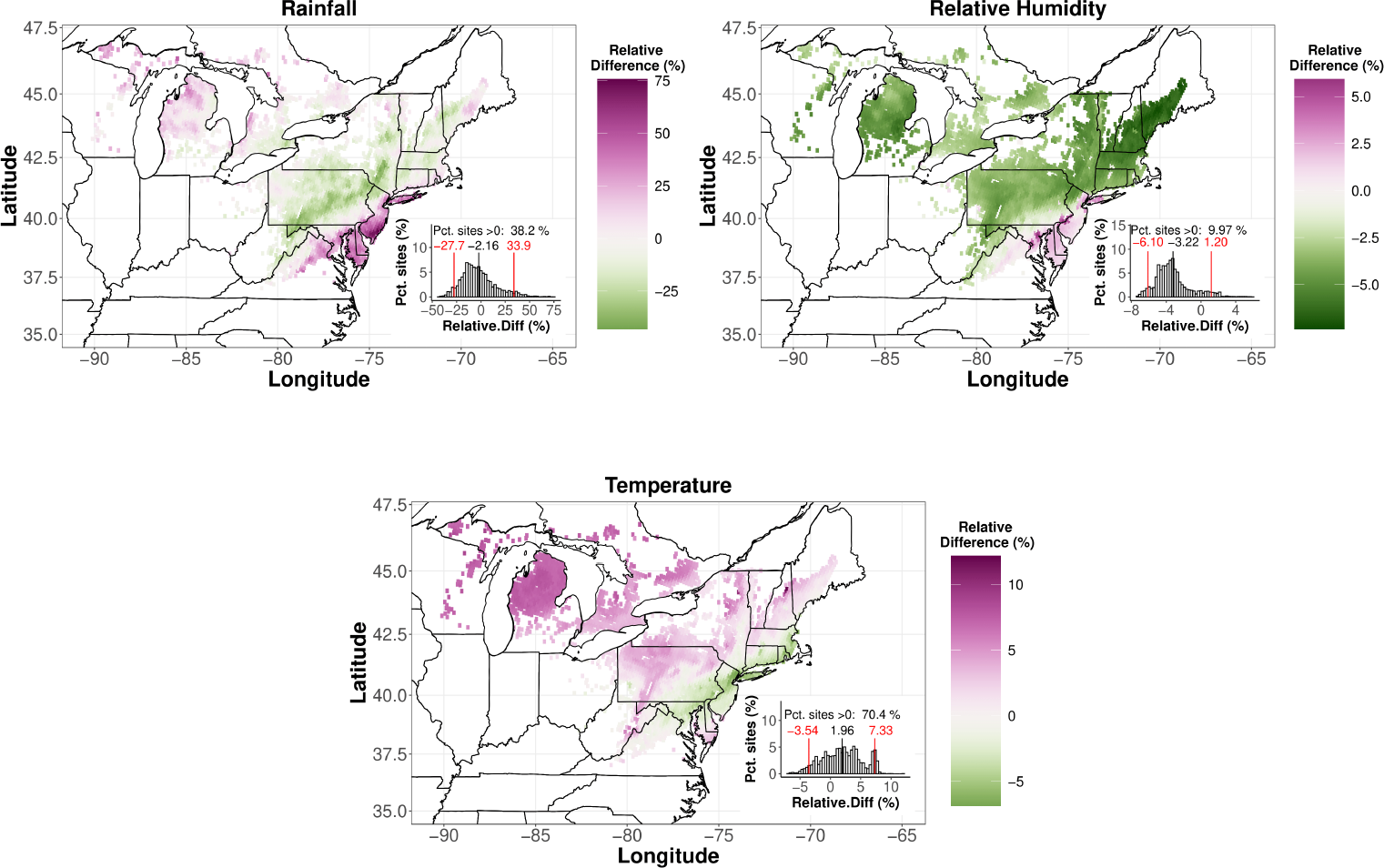
Projected changes due to climate change in key weather variables, in terms of the relative percent change in each variable. If we define *S*_*i,h*,_ and *S*_*i,f*,_ to be the respective historical and future values of a given variable *S* at location *i*, then the relative percent change at location *i* is 100%*×*(*S*_*i,f*,_ *− S*_*i,h*,_)*/S*_*i,h*,_. Here the historical period is 1995-2004, while the future is 2085-2094, under the Relative Concentration Pathway or “RCP” 8.5 scenario; see Supporting Information for projections using the RCP 4.5 scenario, and for the mid-century period 2045-2054. Inset histograms show distributions across locations, with black vertical lines indicating means, and with red vertical lines indicating 5th and 95th percentiles. For context, we note that the mean values of the changes in rainfall, relative humidity and temperature are respectively -0.13 mm/day, -2.43%, and +0.34 ^*?*,^C. Because relative humidity measures the amount of moisture that the atmosphere contains relative to the amount that it can contain, and because the amount of moisture that the atmosphere can contain rises with temperature, sufficiently sharp increases in temperature can lead to reductions in relative humidity even if rainfall increases [25].

As we described, the projections of our model also depend on the density of spongy moth larvae. Spongy moth densities at the beginning of the larval period are determined not just by how many spongy moths were killed by *E. maimaiga* in the preceding generation, but also by generalist small mammal predators [8], whose densities are in turn determined by densities of oak (*Quercus* spp.) acorns [26]. Allowing for these additional species interactions would have seriously complicated our efforts to project initial spongy moth densities, and so we simplified the problem by making projections across the range of densities at which *E. maimaiga* epizootics have been observed in the past.

Unfortunately, our model projects that *E. maimaiga* infection rates will decline at all spongy moth densities and at almost all locations (fig. 2); moreover, reductions in the infection rate are projected to be the most severe at low and intermediate spongy moth densities (fig. 3). Because each female spongy moth can produce hundreds of eggs, low and intermediate density populations can rapidly turn into high density populations [27], and so high *E. maimaiga* infection rates at low and intermediate densities have long played a key role in keeping spongy moth populations in check [28]. Our model projections thus imply that climate change will greatly reduce the impact of *E. maimaiga* on spongy moth populations.

**Fig. 2.**
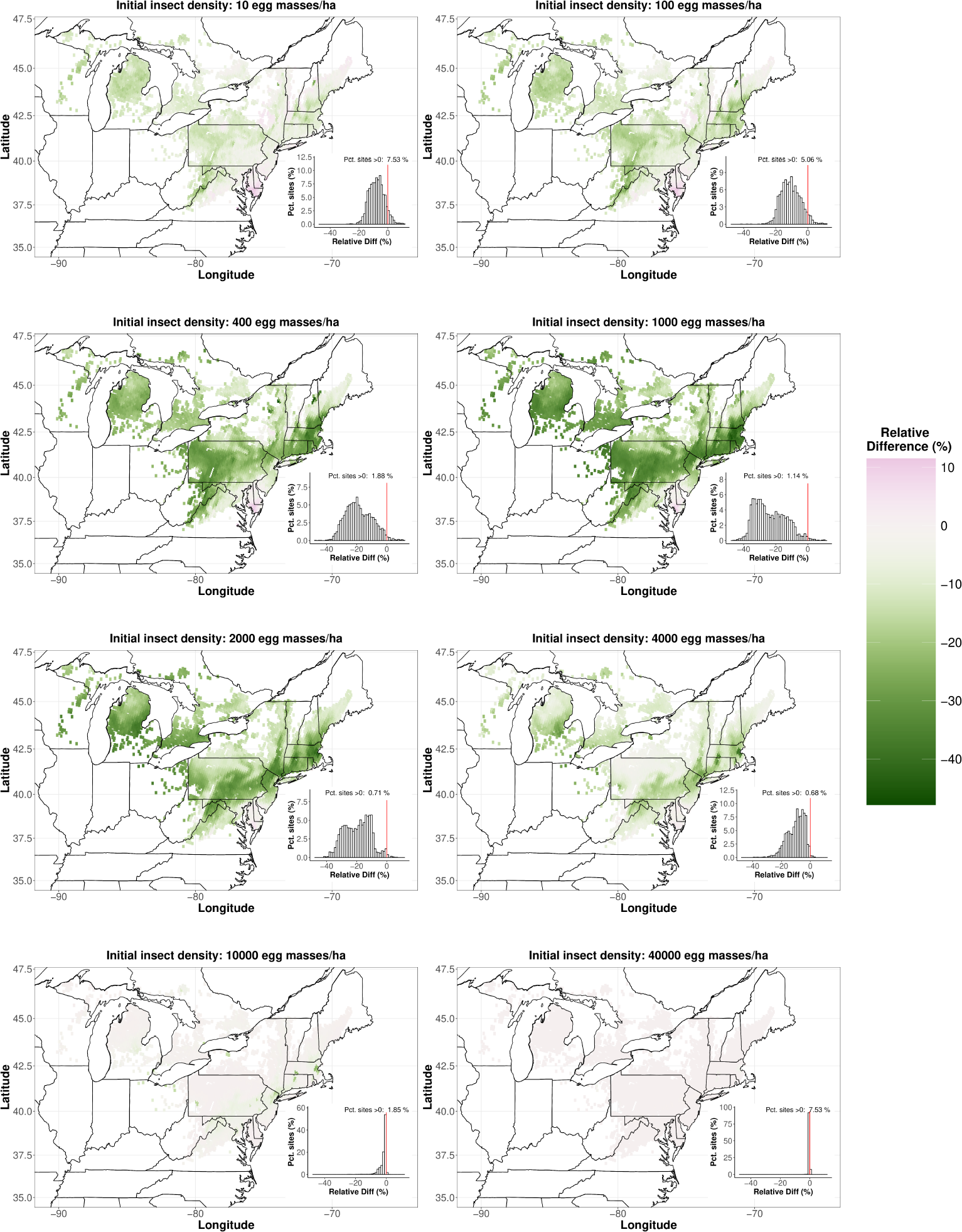
Model projections of changes in the *E. maimaiga* infection rate. Initial spongy moth densities are expressed in units of egg masses/hectare; here we assume 250 eggs per egg mass, a rough average of egg mass sizes in nature [27]. As in fig. 1, we show relative percent changes in the average fraction infected, and we include summary histograms. The red vertical lines in the histograms represent relative changes of 0%.

**Fig. 3.**
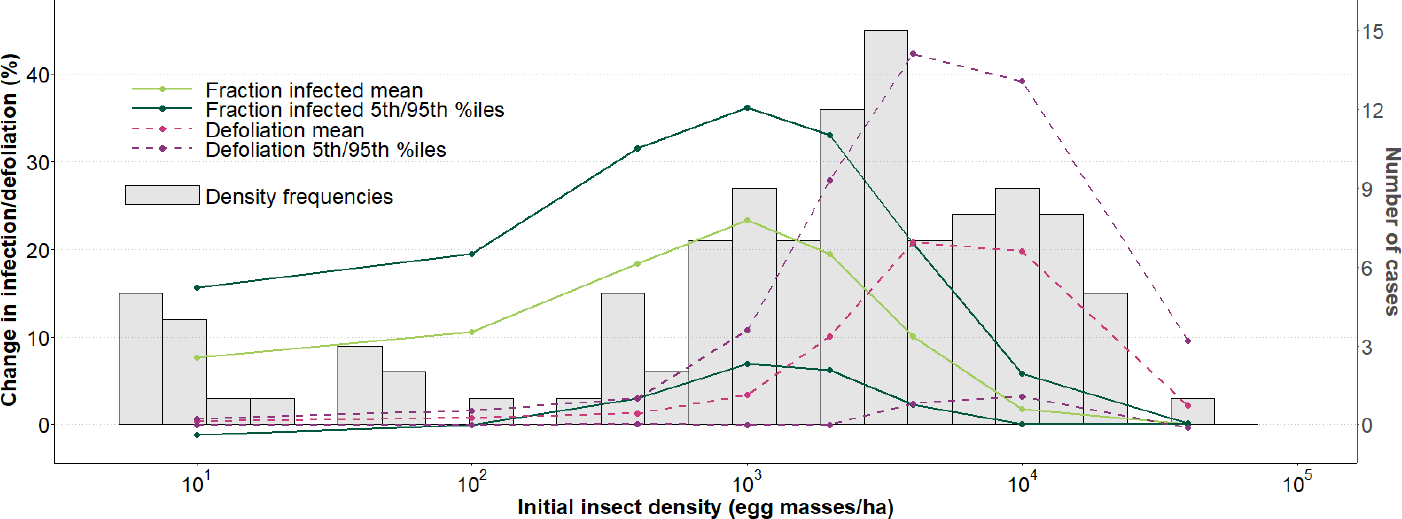
Summary of changes in the fraction infected and in the defoliation rate in the maps in figs. 2 and 4. Because the fraction infected is projected to fall whereas defoliation is projected to rise, here we have multiplied the change in the fraction infected by 1, so that the two types of change can be visualized more easily in the same plot. The light green line then represents the mean relative percent decrease in the fraction infected across sites, while the dark green lines indicate the 5th and 95th percentiles, which were calculated across locations. Small negative values of the lower 5th percentile at low densities indicate that when spongy moth densities are low the fraction infected will increase at a small fraction of locations. The pink dashed line similarly indicates the mean change in the defoliation rate across sites, while the purple dashed lines indicate the corresponding 5th and 95th percentiles, which are again calculated across locations. The gray histograms show the frequency of epizootics at different densities in the literature [20, 28–34]. As a threshold value of what constitutes an epizootic in the data, we used a value of the cumulative fraction infected of 56.7%, which is the model’s mean projected cumulative fraction infected at a density of 1000 egg masses*/*ha under historical weather conditions.

At high spongy moth densities, the model projects that reductions in the *E. maimaiga* infection rate will be more modest, because the negative effects of climate change on *E. maimaiga* will be partly counter-balanced by the positive effects of high spongy moth densities on *E. maimaiga*’s transmission rate. There is nevertheless a range of intermediate to high densities at which *E. maimaiga* epizootics have often occurred and at which the projected reduction in the infection rate is moderate, but at which the projected increase in defoliation is high (figs. 3 and 4). This effect occurs because defoliation rates rise very rapidly with spongy moth densities, and so at intermediate to high densities even modest reductions in the infection rate lead to sharp increases in defoliation in the following year.

**Fig. 4.**
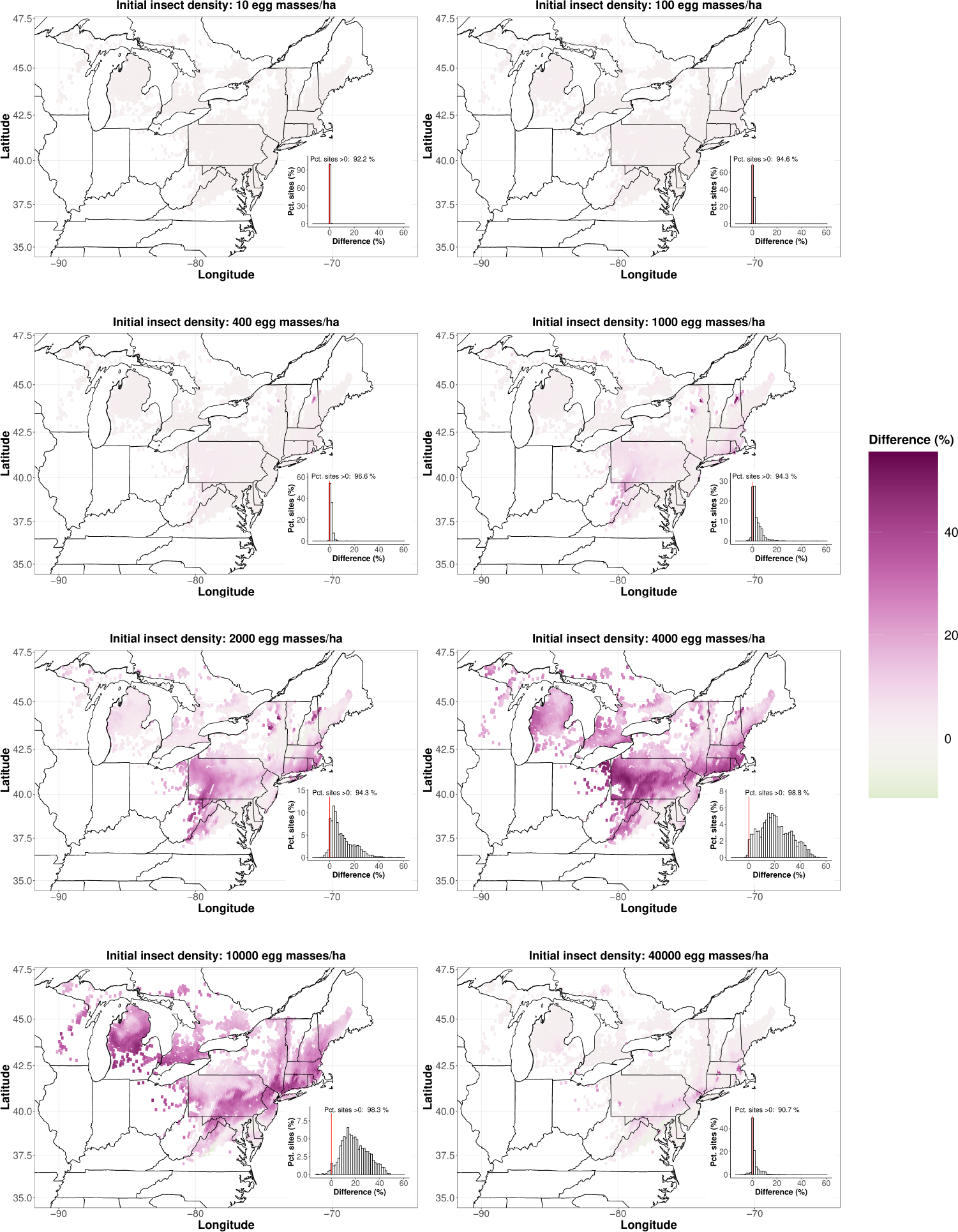
Model projections of changes in spongy moth defoliation levels due to climate change. We calculated post-epizootic defoliation rates by using a nonlinear regression model [16] to convert projected spongy moth densities in the year after the epizootic into percent defoliation. At high density the model accurately projects very high *E. maimaiga* infection rates pre-climate change [20], and so in high density populations defoliation rates are generally very low in the year after an epizootic. These low pre-climate-change defoliation rates can produce extreme projections of relative changes in defoliation, and so here we show untransformed changes instead of relative changes.

Our model thus projects that climate change will have strong negative effects on the *E. maimaiga* infection rate at low to intermediate densities, and strong positive effects on the spongy moth defoliation rate at intermediate to high densities. Climate change will therefore have undesirable effects on the spongy moth across the entire range of densities at which *E. maimaiga* epizootics have been observed in the past, which is essentially the entire range of densities at which spongy moth populations have been observed in the past [27]. Meanwhile, the repeated occurrence of spongy moth outbreaks in the pre-*E. maimaiga*-era means that mortality due to other natural enemies, whether generalist small mammal predators or the spongy moth’s baculovirus pathogen [8], is unlikely to compensate for the decline of *E. maimaiga*.

## Alternative Scenarios

The model that produced the projections in figs. 2-4 assumes that spongy moth larval growth increases linearly with temperature, and that there is no larval mortality due to high temperatures (As we explain in the Supplemental Information, our model implicitly allows for losses that occur when larvae disperse from their egg masses to foliage, which is the most important form of non-disease mortality [35]). Although our model provided a good fit to the data, future temperatures will of course be higher than they were at the time of our study. It is therefore important to ask, would allowing for nonlinear effects of temperature on larval growth and non-disease mortality change our model’s projections?

Because our field data did not show obvious effects of high temperatures on larval growth and non-disease mortality [20], to allow for such effects we estimated the necessary model parameters using laboratory data [36]. Out of concern that the laboratory data provide an unrealistic picture of conditions in the field, we also considered a model in which we sharply but arbitrarily increased the rate at which non-disease mortality increases with temperature relative to the original data.

Although one might expect that allowing for high-temperature effects would lead to at least moderate changes in the projections of our model, instead there was no effect (Supplemental Information). This lack of an effect occurs first because, as we show in the Supplemental Information, future increases in temperature are likely to have only modest effects on spongy moth growth and survival. Second, spongy moth larvae hatch in the spring, and so high temperatures typically do not occur until late in the larval season; meanwhile, the severity of *E. maimaiga* epizootics is determined largely by conditions early in the larval season [20], and so high temperatures late in the season have little effect on *E. maimaiga* infection rates, which therefore typically dwarf non-disease mortality. The ecological effects of climate change on the spongy moth-*E. maimaiga* interaction are thus likely to have a bigger effect on spongy moth defoliation rates than the physiological effects of climate change on the spongy moth.

In the Supplemental Information, we further show that our projections are qualitatively similar across a range of other alternative scenarios, including the RCP 4.5 CO_2,_emissions scenario and scenarios with higher and lower initial densities of overwintering *E. maimaiga* resting spores. Moreover, an analysis of the effects of stochasticity (Supplemental Information) shows that much more severe outcomes are highly likely, whereas much less severe outcomes are not very likely: the mean projections that we show here are therefore optimistic.

## Weather-Only Model

Part of the reason why climate change has such strong effects in our model is that the effects of spongy moth density on transmission in the model translate moderate changes in weather into large changes in *E. maimaiga* infection rates [37]. Previous efforts to quantify the effects of climate on the spongy moth-*E. maimaiga* interaction have in contrast considered the effects of weather but not density [38], and perhaps as a consequence have been only moderately successful at identifying the weather conditions that lead to epizootics. Meanwhile, as we show in fig. 5, a weather-only model projects overly optimistic effects of climate change on the *E. maimaiga* infection rate. A neglect of species interactions can thus cause weather-only approaches to underestimate the negative ecological consequences of climate change.

**Fig. 5.**
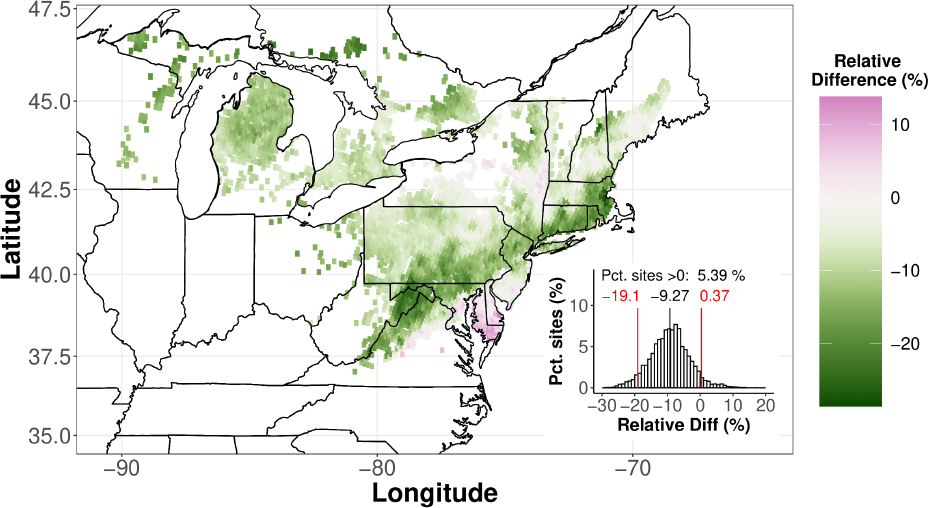
Projections of the weather-only model. Because the *E. maimaiga*-spongy moth interaction is not included in this model, host density has no effect on the model projections, and so we show only a single set of projections.

## Recent Defoliation Data Support the Model’s Projections

Hot, dry conditions in New England in 2015-2018 and in Ontario in 2019-2021 led to the first region-wide spongy moth outbreaks in each region since the advent of *E. maimaiga* in 1989 (fig. 6, see Supporting Information for associated maps) [15, 39]. Meanwhile, as we show in the Supplemental Information, our model projects that increases in defoliation rates over the next 25 years will be nearly as large as the end-of-century projections that we show in fig. 3. The recent New England and Ontario outbreaks thus provide initial confirmation of our model’s projections.

**Fig. 6.**
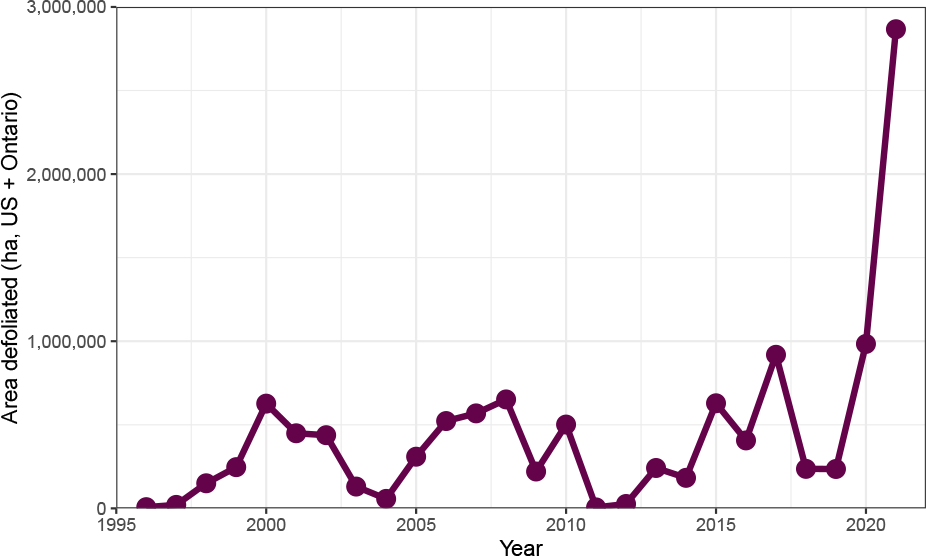
Area defoliated since the widespread introduction of *E. maimaiga* [40].

## Discussion

Our model’s projections are concerning not just because the spongy moth damages economically valuable timber, but also because the spongy moth feeds preferentially on oaks, *Quercus* spp. The spongy moth has therefore been an important factor driving “oak decline” [10], the ongoing replacement of oaks by maples, *Acer* spp., in North America [41]. Compared to oaks, maples provide fewer resources for native insects, songbirds and small mammals [42]; the effects of climate change on the *E. maimaiga*-spongy moth interaction thus provide an important example of how the effects of climate change can radiate through natural communities.

Our results are also relevant to projections of how climate change will affect hard-wood forests. Models of the effects of climate change on tree growth have projected that increasing CO_2,_ concentrations may have fertilizing effects on forests [43], especially if trees can acclimate to drought [44]. Tree-growth models, however, focus on the effects of climate change on tree physiology, and therefore either neglect insects entirely, or include insects simply as disturbances [45]. By explicitly tracking insect densities, our model instead demonstrates that the effects of climate change on insect pathogens may result in more severely negative effects of climate change on forests than anticipated. Whether or not spongy moth defoliation will therefore counter-balance the fertilizing effects of CO_2,_ is as yet unknown, however, and so an important future direction is to unite the ecological approach that we have taken here with the physiological approach of tree-growth models.

An additional future direction is to allow for effects on the spongy moth of climate-change driven alterations in the density of oaks versus other host trees, a mechanism that has been shown to have detectable effects on spongy moth defoliation [46]. Also, the spongy moth’s use of multiple host trees means that phenological mismatch with tree budburst usually has only small effects on the insect’s population growth rate [35], but allowing for multiple tree species in our model would make it possible to test whether such effects would be strengthened by climate change.

Our work provides strong support for the growing consensus that climate change may negatively affect species interactions that are important to humans [47]. The negative consequences of climate change for human societies may thus be more severe than previously believed.

## Supplementary information

This article includes a Supplementary information file.

## Acknowledgments

We thank NSF OPUS grant 2043796, SERDP DOE contract DE-AC0206CH11357, and U. Chicago RCC. J.L. thanks the Chinese Scholarship Council. A. Liebhold provided defoliation data and useful advice.

