## Supplemental Information for "Climate-Change Driven Decline of an Insect Pathogen Increases the Risk of Defoliation by a Forest Pest Insect"

**6 Effects of Stochasticity on the Model Projections** **29**

**7 Comparison of Recent and Historical Defoliation Maps** **33**

### 1 Model Structure

#### 1.1 SEIR Model

##### 1.1.1 Overview

As we mentioned in the main text, in previous work we tested our ecological model using a data set that we collected over three years on a 300-km-long north-south transect in the lower peninsula of Michigan [1]. That data set includes measurements of both *E. maimaiga* infection rates and weather variables, specifically temperature, rainfall, and relative humidity, in each of seven study plots.

*E. maimaiga* infection rates at some sites in some years were determined mostly by spongy moth densities, while at other sites and in other years infection rates were determined mostly by weather conditions. For example, of two plots with densities of about 240 egg masses per hectare, one plot experienced low rainfall while the other experienced moderate rainfall, yet both had low weekly infection rates. Density thus played a key role in determining the infection rates in the two plots. Meanwhile, in a plot at a density of 2300 egg masses per hectare, high rainfall and high relative humidity produced weekly infection rates that were often higher than the weekly infection rates in a plot that had a density of 10,000 egg masses per hectare but which experienced only moderate rainfall and moderate humidity. Our data thus reflected the effects of both spongy moth density and weather, and therefore allowed us to disentangle the effects of the two different factors on *E. maimaiga* infection rates.

Ultimately, however, variation in weather and variation in spongy moth density together were not enough to fully explain the data, and so we also allowed for variation in resting spore densities. To do this, we included a set of parameters that separately represented the resting spore density in each study plot, and we estimated each of these resting spore densities from the data. Although in principle it would have been possible to directly estimate resting spore densities by using a microscope to count spores in soil samples [2], our indirect estimation procedure allowed us to avoid making assumptions about where resting spores were located in the environment. This is important because larvae that produce resting spores die at or near the base of trees, and so resting spores may overwinter either on bark or in the soil [3].

Because resting spore variation also did not allow the model to fully explain the data, we further included stochasticity, to allow for variation in the *E. maimaiga* transmission rate above and beyond the effects of spongy moth density, resting spore density and weather. The best model was then able to explain almost all of the variation in the data.

As we described, part of the reason why we were able to separately estimate the effects of three different weather variables, spongy moth densities, and resting spore densities is because the weather variables, the spongy moth densities and the *E. maimaiga* infection rates all varied greatly among study sites. Two additional reasons, however, are first that we fit our models to time series of *E. maimaiga* infection rates, and so we were able to explicitly take into account fluctuations in infection rates that resulted from changes in weather conditions, changes in spongy moth densities, and changes in conidia densities during epizootics.

A second reason why we were able to separately estimate the effects of weather and densities is that we deployed experimental cages of lab-reared larvae in each plot, which we used to quantify infection rates over successive 24-hour periods. Because of the short deployment times of the cages, the infection rate among the caged larvae provided snapshots of *E. maimaiga* transmission in the near-absence of changes in conidia densities. Because the resulting data allowed only for the effects of *E. maimaiga* transmission, they made it possible to estimate the transmission parameters with a greater degree of independence from the other model parameters.

Previous studies of *E. maimaiga* epizootics have in contrast included only observational data, and have almost invariably quantified infection rates either at a single time point, or by condensing time series of infection rates from an epizootic into a single estimate of the cumulative infection rate for that epizootic [4–10]. These approaches make it possible to use logistic regression models to analyze epizootic data, but they make it difficult to test more than very simple hypotheses about the mechanisms driving epizootics. Because we instead fit our models to time series of infection rates, our approach allowed us to test more complex hypotheses; as we explained in further detail in our previous study [1], by using a state-of-the-art nonlinear fitting routine we also maintained a high level of statistical robustness.

##### 1.1.2 Model Equations

Here we briefly summarize the structure of our SEIR-type model: see Kyle et al. [1] for more details. The model consists of random ordinary differential equations [11] numerically integrated on a daily time scale:

$$\frac{dS_\tau}{dt} = -\nu_{R,\tau} S_\tau R_\tau - \nu_{C,\tau} S_\tau C_\tau, \tag{1}$$

$$\frac{dE_{\tau,1}}{dt} = \nu_{R,\tau} S_\tau R_\tau + \nu_{C,\tau} S_\tau C_\tau - m\lambda E_{\tau,1}, \tag{2}$$

$$\frac{dE_{\tau,j}}{dt} = m\lambda E_{\tau,j-1} - m\lambda E_{\tau,j}, \qquad j = 2, ..., m. \tag{3}$$

$$\frac{dC_\tau}{dt} = m\lambda E_{\tau,m} - \mu_{C,\tau} C_\tau. \tag{4}$$

Here $S_\tau$, $R_\tau$, $E_{\tau,j}$ and $C_\tau$ are the densities of susceptible hosts, resting spores, exposed larvae in class $j$, and conidia on day $\tau$. We then integrate from time $t = 0$ to time $t = 1$ on each day $\tau$, with the initial value of each state variable set equal to its value at the end of the preceding day.

Resting spore density was set to the initial value $R(0)$ for $T_g \leq \tau \leq T_e$, and to zero for $\tau < T_g$ and $\tau > T_e$, where the beginning $T_g$ and end $T_e$ of the resting spore germination period were estimated from our data. We then varied the initial density of hosts $S_0(0)$ and resting spores $R(0)$ across scenarios.

On day $\tau$, the conidia decay rate $\mu_{C,\tau}$ and the transmission rates $\nu_{R,\tau}$ for resting spores $R_\tau$, and $\nu_{C,\tau}$ for conidia $C_\tau$ are determined by stochasticity and by the weather projected for that day by our climate-change model [12]. Larval infection risk increases as larvae increase in size, as determined by a degree-day dependent function $D(\tau)$ that we describe in the next subsection.

To incorporate weather into the model we used information from the literature to construct functions that determine how conidia transmission $\nu_{C,\tau}$, resting spore transmission $\nu_{R,\tau}$ and conidia decay $\mu_{C,\tau}$ depend on weather, and we fit the parameters of each function to our epizootic data, using our daily measurements of temperature, relative humidity, and rainfall as covariates [1].

The transmission function for resting spores is a saturating function of accumulated daily rainfall over the previous 10 days $p(\tau)$:

$$\nu_{R,\tau} = D(\tau)\left(\frac{\psi_1}{1+\psi_2\exp(-\psi_3 p(\tau))} - \frac{\psi_1}{\psi_2+1}\right)\exp(\epsilon_{R,\tau}). \tag{5}$$

This function was constructed to ensure that resting spore transmission is zero when cumulative rainfall $p(\tau) = 0$. The parameters $\psi_1$-$\psi_3$ were estimated from the data, as was the standard deviation of the normally distributed, zero-mean stochastic term $\epsilon_{R,\tau}$ [1].

The conidia transmission function $\nu_{C,\tau}$ increases exponentially with minimum daily relative humidity $m(\tau)$;

$$\nu_{C,\tau} = D(\tau)\psi_4\exp\big(\psi_5 m(\tau)\big)\exp(\epsilon_{C,\tau}). \tag{6}$$

As in equation (5), the parameters $\psi_4$ and $\psi_5$ were estimated from the data, as was the standard deviation of the zero-mean, normally distributed stochastic term $\epsilon_{C,\tau}$ [1].

Because high temperatures inhibit conidia survival [13], we assumed that conidia decay $\mu_{C,\tau}$ is an exponentially increasing function of daily maximum temperature, so that conidia survival falls with temperature;

$$\mu_{C,\tau} = \psi_6\exp\big(\psi_7 h(\tau)\big). \tag{7}$$

Here $h(\tau)$ is the maximum temperature on day $\tau$, and the parameters $\psi_6$ and $\psi_7$ were estimated from the data [1].

To construct the weather-only version of the SEIR model, we assumed that *E. maimaiga* transmission is unaffected by the density of conidia or resting spores. The weather-only model therefore tracks only the density of susceptible larvae $S$ and the density of larvae in exposure class $E_j$, according to:

$$\frac{dS_\tau}{dt} = -\nu_{F,\tau}S_\tau. \tag{8}$$

$$\frac{dE_{\tau,1}}{dt} = \nu_{F,\tau}S_\tau - m\lambda E_{\tau,1}, \tag{9}$$

$$\frac{dE_{\tau,j}}{dt} = m\lambda E_{\tau,j-1} - m\lambda E_{\tau,j}, \qquad j = 2, ..., m. \tag{10}$$

Here $\nu_{F,\tau}$ depends on temperature $h(\tau)$ and rainfall $p(\tau)$;

$$\nu_{F,\tau} = D(\tau)\left(\frac{\psi_8}{1+\psi_9\exp(-\psi_{10}p(\tau))} - \frac{\psi_8}{\psi_9+1}\right)$$

$$\times \exp\Big(-\psi_{11} h(\tau) \exp(\epsilon_{F,\tau})\Big). \quad (11)$$

As in the rain-dependent transmission function in the eco-climate model, transmission $\nu_{F,\tau}$ is 0 when no rain occurs. The parameters $\psi_8$-$\psi_{11}$ and the standard deviation of the mean-zero stochasticity term $\epsilon_{F,\tau}$ were estimated from the data [1]. Stochasticity $\epsilon_{F,\tau}$ is here parameterized differently than in the full eco-climate model, due to an oversight during the model-fitting process. Re-fitting the full model using the parameterization in equation (11) confirmed that this difference affects only the scaling of the standard deviation in stochasticity, and thus has no effect on our results.

##### 1.2 Effects of Temperature on Hatch Time and Larval Growth

Because *L. dispar* eggs hatch in response to an accumulation of degree days [14], the timing of spongy moth larval emergence in the model depends on temperature [1]. Larval emergence in the model therefore occurs earlier in the year as temperatures increase due to climate change.

As is typical of insects [15], spongy moth larval growth increases with temperature [16], and larval size in the model therefore also increases with temperature. Larval size is important in the model first because, as in nature, epizootics in the model end when larvae have completed their development, and, again as in nature, larger larvae in the model are at greater risk of *E. maimaiga* infection [17]. This latter effect is so strong that when we left it out of an early version of the model that version could not reproduce the sharp increases in the infection rate that often occur late in the larval season [1].

In laboratory experiments, constant temperatures above 28-30° C can cause larval growth rates to plateau or even decline [18]. When temperatures instead vary between daytime and nighttime, however, larval growth rates instead depend in a complex way on the length of time over which temperatures vary, as we describe below [19]. Because of these complexities, and because the effects of high temperatures on growth rates are not very strong, it is unlikely that we would have been able to accurately estimate nonlinear effects of temperature on larval growth rates from our epizootic data. Moreover, the model that uses a linear growth rate function gave a good fit to the data, and so it is likely that allowing for nonlinear effects of temperature would have led to over-parameterization of the model and thus to unreliable model projections.

To maintain consistent use of field data for estimating parameters, in the main text we include results only for a model for which larval growth increases linearly with temperature, and for which larval mortality is due only to *E. maimaiga* infection. (Because in our field study we quantified spongy moth densities before larvae had hatched from their egg masses, whereas we quantified infection rates after hatched larvae had dispersed from their egg masses onto foliage, our model's parameters implicitly take into account the non-disease mortality that occurs as larvae disperse from their egg masses, which is by far the most important source of non-disease mortality [20]). In what follows, however, we also include results for a model in which larval growth saturates at high temperatures, and in which non-disease larval mortality increases at high temperatures, with the rates of decline or increase estimated from laboratory data. As we describe below, the resulting model gives results that are effectively identical to the results that we present in the main text.

### 2 Using Literature Data To Test the Eco-Climate Model

#### 2.1 High Rainfall and High Resting Spores Can Explain High Infection Rates at Low Densities

Although in our previous study our model provided an excellent fit to our data [1], that does not necessarily mean that the model can explain data sets collected in previous studies. Unfortunately, however, as we mentioned in the main text previous studies did not collect the daily local weather data that our model needs to make projections. Moreover, as we mentioned in the previous section, previous studies collected only single time points, or condensed time series data into summary statistics of the data. It is therefore not possible to carry out the kind of statistically robust comparison of the model to literature data that we carried out in our previous study. It is nevertheless possible to at least qualitatively compare the model to literature data, and we therefore do so here.

First, multiple studies have documented *E. maimaiga* infection rates of more than 50% at spongy moth densities of 400 or more egg masses per hectare [6–8, 21, 22]. For low to moderate resting spore densities and average weather conditions, our model similarly shows infection rates of more than 50% at densities of 400 or more egg masses per hectare, as we showed in our previous work [1]. Moreover, the infection rate in our model increases with spongy moth density at a roughly similar rate as in the work of Hajek and van Nouhuys [8], who presented *E. maimaiga* infection rate data collected at a wide range of densities and locations.

It is also true, however, that *E. maimaiga* infection rates sometimes exceed 50% even in low density spongy moth populations. As we will discuss, such cases often occur in years of high rainfall or within a few years after a spongy moth population has crashed because of an *E. maimaiga* epizootic; because in our previous work we studied only populations that were in the midst of crashing, we did not observe any of the latter cases. Here we nevertheless show that our model can account for observations of high infection rates in low density populations by invoking high densities of resting spores and/or high rainfall.

For context, we first quantify the frequency with which high mortality has been reported in low-density spongy moth populations in the literature. Table 1 shows data from what is to our knowledge every case for which *E. maimaiga* mortality was estimated in a spongy moth population with a reported density of about 100 egg masses per hectare or less (We do not include studies that instead used proxies for density, on the grounds that the relationship between the proxies and density is unclear). As the table shows, in 16 of 39 cases (counting each three-year period for the Hajek and van Nohouys [8] data set as three cases each), mortality was above 0.5, while in 6 of 39 it was above 0.85. High *E. maimaiga* mortality has thus been observed

| Citation | Collection Year/Plot | EM/ha ± SE | Fract. *E.m.* |
|---|---|---|---|
| Hajek et al. 1990 [4] | 1989 A | 45.3 ± 15.3 | 0.877 |
| | 1989 B | 42.5 ± 11.6 | 0.782 |
| | 1989 C | 42.5 ± 9.7 | 0.786 |
| | 1989 D | 19.8 ± 7.1 | 0.603 |
| Hajek et al. 1996 [5] | 1994 | 0 | 0.167 |
| | 1994 | 0 | 0.636 |
| | 1994 | 0 | 0.129 |
| Hajek et al. 1997 [6] | 1993 | 103 ± 22 | 0.186 |
| | 1994 | 0 | 0.70 |
| | 1995 | 0 | 0.06 |
| | 1996 | 11 ± 11 | 0.946 |
| Liebhold et al. 2013 [21] | 2007 or 2008 | 10 | 0.902 |
| | 2007 or 2008 | 10 | 0.950 |
| | 2007 or 2008 | 10 | 0.616 |
| | 2007 or 2008 | 10 | 0.987 |
| | 2007 or 2008 | 50 | 0.846 |
| Hajek & van Nouhuys 2016 [8] | | | |
| Stable | 1996-1998 | 25.9 | 0 |
| | 1999-2001 | 22.2 | 0.177 |
| | 2002-2004 | 18.5 | 0.141 |
| | 2005-2007 | 7.41 | 0.147 |
| | 2008-2010 | 11.1 | 0.110 |
| Outbreak-prone | 2009 | 0 | 0.124 |
| | 2009 | 0 | 0.274 |
| | 2009 | 0 | 0.335 |
| | 2009 | 0 | 0.351 |
| | 2009 | 0 | 0.595 |
| | 2009 | 0 | 0.667 |
| | 2009 | 49 | 0.697 |
| | 2009 | 0 | 0.982 |

**Table 1** Literature data showing *E. maimaiga* mortality at spongy moth densities of 100 egg masses per hectare or less. The data from Hajek and van Nouhuys [8] for the "Stable" sites are averages over the three years in each period. Note that sites with 0 egg masses per hectare of course had at least some spongy moth larvae, because otherwise it would have been impossible to estimate the *E. maimaiga* infection rate. Moreover, estimating egg mass densities that are below 100 egg masses per hectare is extremely difficult, and so observations of 0 egg masses may in some cases represent substantially higher densities. Finally, the years for the Liebhold et al. [21] data include 2007-2009, but the data associating sites and years are no longer available. Comparison of the Liebhold et al. and the Hajek & van Nouhuys data, however, made clear that the data from the "Outbreak-prone region" in Hajek and van Nouhuys included the 2009 data from Liebhold et al. but not the 2007 or 2008 data. We were thus able to rule out 2009 as the collection year for the Liebhold et al. low-density data.

in low density spongy moth populations at least moderately often, and so here we consider the extent to which our model can reproduce these observations.

Notably, Hajek and colleagues [4, 6, 8] have argued that cases of high infection rates at low densities in nature have generally resulted from high rainfall and/or the carryover of high resting spore densities from preceding generations. In testing whether our model can reproduce cases of high infection rates at low densities, we therefore tested whether incorporating high rainfall, high resting spore densities or both into our model produces model projections that are close to the data.

As we will show, the main challenge to the model comes from the cases in Table 1 in which the fractional *E. maimaiga* mortality was greater than 0.9. This includes the three sites from Liebhold et al. [21] at 10 egg masses per hectare, the single site from Hajek [6] at 11 egg masses per hectare, and the single site from Hajek and van Nouhuys [8] at a reported density of 0 egg masses per hectare. Because it is of course true that there can be no epizootic if there are no larvae, in what follows we treat the site from Hajek and van Nouhuys as though it were in fact 10 egg masses/hectare. We use 10 egg masses/ha as an intermediate value on a $\log_{10}$ scale between the reported value of 0 and the likely upper limit of 100 egg masses/ha. Meanwhile, differences in model infection rates for densities of 10 versus 11 egg masses/hectare are too small to be of much interest, and so in what follows we similarly treat the 11 egg mass/ha site as if it had a density of 10 egg masses per hectare.

To compare our model to the data, it is of course important that we take into account measurement error, especially because the difficulty of locating larvae in low-density spongy moth populations means that the studies in question did not have particularly large sample sizes. Because most of the studies did not provide estimates of the error in their measurements of spongy moth densities, here we focus on error in their measurements of *E. maimaiga* infection rates. For the Liebhold et al. [21] study in particular, a further complication is that the infection rate was calculated as a "proportional hazard", meaning the mortality rate that *E. maimaiga* would have produced if there were no other competing sources of mortality. Because the original calculations are no longer available, we cannot allow for this additional error, and we thus do not attempt to explain the proportional-hazard calculation, other than to say that it almost certainly increased the measurement error.

The simplest way to allow for measurement error in the infection rates is therefore to add measurement error to our model's projections, thereby following a standard procedure for this type of data [23]. Because the Liebhold et al. [21] data provide the largest number of cases of very high *E. maimaiga* mortality at very low spongy moth densities, and because the Hajek and Elkinton [4] and Hajek and van Nouhuys [8] studies provide few details of their sampling schemes, in allowing for measurement error we followed the sampling schemes in Liebhold et al.

The missing data from Liebhold et al. [21] include the data associating years with study sites; as we explain in the legend for Table 1, however, we at least know that the cases of high *E. maimaiga* mortality came from data collected in either 2007 or 2008. We then used the sampling schemes for both 2007 and 2008, but the two sampling schemes gave very similar results, and so here we present results only for the 2007 sampling scheme.

In that year, Liebhold et al. estimated cumulative mortality by making five collections of larvae at intervals of 4 days, sampling 50 larvae on each collection date. To determine the survival rate during each time period, Liebhold et al. then reared the collected larvae until the next collection date, or for 30 days in the case of the last larval collection. Cumulative mortality was then again calculated as one minus the product of survival rates across time periods; a similar procedure was apparently used in the other studies, except that Hajek and Elkinton [4] estimated infection rates only for single weeks.

Liebhold et al. [21] reported that their data were based on collections of late-instar larvae, as were the data from the outbreak-prone areas in Hajek and van Nouhuys [8], and the single-time-point data in Hajek and Elkinton [4]. We therefore assumed that collections started from the fifth week after larval hatch, and thus at least a few days after resting spores had begun to germinate. In our previous work, the sampling error for *E. maimaiga* infection rates was well described by a beta-binomial distribution, and we therefore use that distribution here, along with the variance that we estimated for that distribution [1]. We then inserted the reported sample size (50 larvae), our estimate of the sampling variance, and our model's projection of the infection rate into a beta-binomial distribution, and we randomly drew the number of surviving larvae from this distribution. This procedure produced a stochastic realization of the fraction infected that took into account the error introduced by the sampling process. As in the data, the cumulative mortality in the model was calculated as one minus the product of the survival rate in each period.

Because we do not have detailed local weather data for any of the study sites in the literature, we used a Richardson weather generator to generate values of the weather covariates that we needed to generate model projections (Note that our far more realistic climate-change model cannot easily allow for user-generated increases in rainfall, and its projections are therefore inappropriate for the test that we carry out here). Richardson weather generators consist of sets of linear models, which in our case were used to express relative humidity and temperature as functions of rainfall; in our previous study, we fit the parameters of our generator to the weather data that we collected [1]. Our weather generator therefore allows for realistic correlations between weather covariates. As input, the generator uses the daily probability of rainfall, the average amount of rainfall and the variance in the amount of rainfall across all days for which there was measurable rainfall.

We then first considered a moderate rainfall case, which means that we used the mean, the variance and the probability of daily rainfall from our previous work [1]. As we show in fig. 8 in the next section, the weather in our previous study encompassed most of the variation in weather across the range of the spongy moth in eastern North America, and so it seemed likely that the mean and variance of the rainfall in our previous study would lead to realistic weather conditions at other locations, including the locations in the studies in Table 1. We then carried out additional simulations in which we allowed for higher rainfall by multiplying the mean rainfall by 3, and by increasing the daily probability of rainfall from 0.4 to 0.8. To ensure that rainfall was not always high, in the high-rainfall case we multiplied the variance by 2. By increasing the rainfall levels in the model we thus allowed for the high rainfall levels reported in Hajek and Elkinton [4]. Liebhold et al. [21] and Hajek and van Nouhuys [8] in contrast provide no information about weather conditions during their studies.

To understand the model's results, it is important to remember that the model is stochastic and that we are allowing for measurement error; in our tests of the model, we therefore conclude that the model can reproduce the data if the upper 95th percentile of the model projections, including measurement error, includes the data. Fig. 1 then shows that the model can indeed explain the data. For moderate rainfall and a resting spore density of 0.05 resting spores/$m^2$, the model's upper 95th percentile exceeds a fractional mortality of 0.9, while at 0.1 resting spores/m$^2$ the upper 95th percentile exceeds or matches the mortality values for the remaining three cases in Table 1 for which mortality was above 0.9. These results hold for both 10 egg masses per hectare and 50 egg masses per hectare.

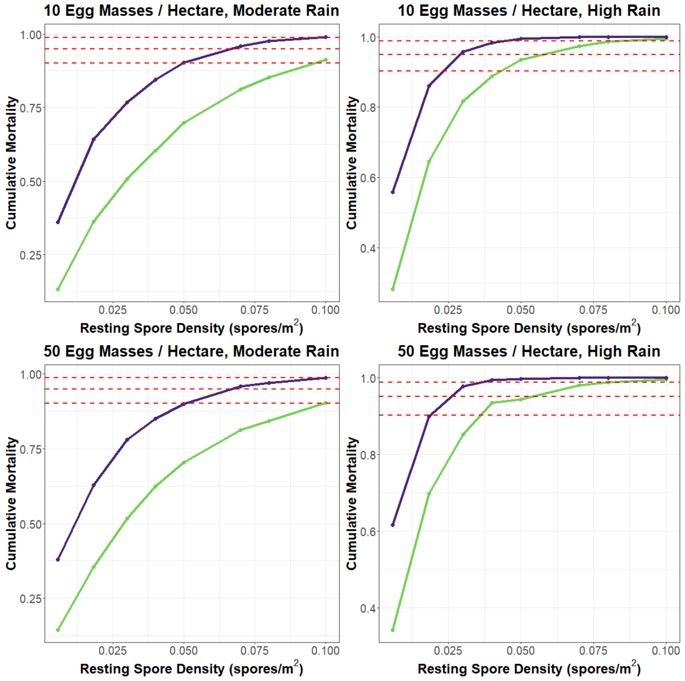

**Fig. 1** Effects of resting spore density on *E. maimaiga* mortality. The curved lines indicate the means (lower, light green) and the 95th percentiles (upper, dark blue) of model projections. The dashed, red horizontal lines indicate the cases of mortality above 0.9 from Table 1; note that the highest mortality from [6] and the second-highest mortality at 10 egg masses per hectare from [21] are both nearly 0.95, and are therefore represented by a single line. To quantify model percentiles, we first generated 100 model realizations, each including mortality rates over the 5 collection periods, and then we generated 10 values of the number of observed larvae killed out of 50 for each time period in each realization. By permuting these latter sets of samples, we generated $10^5$ total sample realizations for each of the 100 model projections, for a total of $10^7$ model realizations. The overlap between the upper 95th percentiles and the data shows that increasing the rainfall or the resting spore density in the model allows the model to reproduce the data.

Fig. 1 further shows that, if we allow for higher rainfall, then the resting spore densities that we need to explain the data are substantially lower, especially if the spongy moth density is somewhat higher. At 10 egg masses per hectare, a resting spore density of 0.03 per m$^2$ is sufficient to explain the three lowest of the four cases in question, while a density of 0.05 resting spores per m$^2$ can explain the remaining case. When the spongy moth density is 50 egg masses per hectare, then the highest estimate of resting-spore density in our previous study, 0.0183 spores per m$^2$, is sufficient for the model to explain the lowest of the four cases, and the model can explain the remaining cases at 0.03 resting spores per $m^2$. The figure further shows that at increased rainfall and a spongy moth density of 50 egg masses per hectare, resting spore densities between the intermediate estimate of 0.00571 that we use in the main text and our highest estimate of 0.0183 can produce mortality rates between 0.6 and 0.9, and thus can explain many of the cases of high mortality in Table 1. It is also worth re-stating that our estimates of the measurement error in the data are necessarily lower bounds, and so the actual measurement error may be considerably larger than the error that we have included here.

Our model can thus explain cases in which the fraction infected is above 0.9 at low egg mass densities by invoking high resting spore densities or high rainfall. We therefore argue that our model has survived a qualitative test with literature data. Moreover, as we mentioned high resting spore densities and high rainfall have similarly been invoked by Hajek and colleagues, and so our model's explanation for the data is supported by previous authors.

As we mentioned earlier, part of the reason why the data to which we fit our model did not include cases of very low spongy moth densities is because we focused on populations that were in the early stages of collapsing, and that therefore had low resting spore densities [1]. The data from low-density sites in Table 1 in contrast were in at least some cases from populations that were known to have already collapsed, while for other sites the recent history of the population was not reported, and so those sites may also have already collapsed. Because our concern here is with whether *E. maimaiga* can cause high mortality in rising populations as opposed to whether *E. maimaiga* can cause high mortality in collapsed populations, in our projections we use estimates of resting-spore densities from our previous study.

#### 2.2 Spatial Scale and Conidia Dispersal in the Data and in the Model

Besides high resting spore densities and high rainfall, another possible explanation for cases of high *E. maimaiga* mortality at low spongy moth densities is the dispersal of infectious conidia between high density and low density study sites. Conidia are wind borne, and can therefore travel distances of 10-100 kilometers in a single season [24, 25]. Meanwhile, as fig. 2 shows, in the Liebhold et al. study [21] seven of the pairwise distances between study sites were less than 10 km, and the average pairwise distance between sites was less than 30 km, while in our previous study [1] the distances between sites ranged from 85 to 350 km (fig. 2), and in the study of Hajek and van Nouhuys [8] sites were scattered across Virginia, West Virginia, Pennsylvania and Maryland, and thus were often far apart. Indeed, in the Liebhold et al. study the small distances between sites may explain why it was not possible to detect an effect of spongy moth density on infection rates, whereas in both our study and in the Hajek and van Nouhuys study it was possible to detect an effect of spongy moth density on infection rates. Unfortunately, however, the data file associating spongy moth densities with study plot locations for the Liebhold et al. study is no longer available, so we cannot directly test the extent to which the movement of conidia affected infection rates in that study.

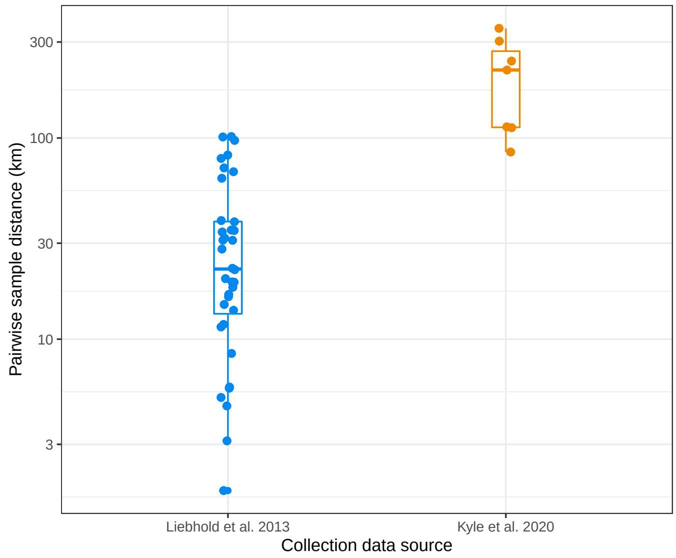

**Fig. 2** Pairwise distances between study sites in the study of Liebhold et al. [21] and in our previous study [1]. Points show the data, while the boxes show the upper 75th and lower 25th percentiles and the vertical lines show two standard errors of the mean. The differences in the distances between plots in the two studies are large enough that distances are shown on a $\log_{10}$ scale.

Issues of conidia dispersal and spatial scale are also relevant to the construction of our model. The model makes projections in 12km × 12km squares, thereby allowing for high levels of conidia dispersal at scales of up to about 17 kilometers, the approximate length of the diagonals of our model grid squares. The model thus allows for the possibility that conidia dispersal can average out infection rates at small to medium spatial scales. The extent of long-distance dispersal at larger spatial scales, however, is not sufficiently well known to allow us to estimate long-distance dispersal rates, and so we did not include long-distance dispersal in the model. Moreover, one of our goals is to determine the extent to which local weather can alter *E. maimaiga* infection rates; because we allow only for medium-distance dispersal, our approach allows for the possibility that *E. maimaiga* infection rates will be reduced by locally unfavorable climate change. Our model structure thus ensures that the effects of climate change in the model are not accidentally eliminated by the unwitting use of over-estimates of conidia dispersal rates.

### 3 Creating Maps

#### 3.1 Generating Model Projections Across Space

Because our models are stochastic, our projections consist of averages over model realizations. At each map location in each year we thus averaged the fraction infected over 100 realizations or "runs" of the model, to produce an average at that location in that year. Next, we averaged the value at each map location across the years 1995-2004, to produce an historical average for each map location. We then similarly averaged the value at each map location across the the years 2085-2094 (main text) or 2045-2054 (below), to produce a future average for each map location.

#### 3.2 Identifying Locations That Are At Risk of Defoliation

Spongy moth defoliation generally occurs in forests that are dominated by oaks (*Quercus* spp.), the preferred food source of spongy moth larvae [14]. Although the distribution of oak-dominated forests may be altered by climate change, it is not clear how soon such alterations will occur [26]. Accounting for possible shifts in the location of oak-dominated forests would thus have seriously complicated our efforts to make projections. We therefore used areas of previous defoliation to determine where defoliation will likely occur in the future, and thus to determine where we should make projections.

To identify areas of previous defoliation, we used two spatially referenced data sets. The first data set, provided by Dr. Sandy Liebhold of the US Forest Service, maps defoliation in the USA from 1975 to 2019 [27]. The data from 1975-2000 were originally from hand-drawn maps that Liebhold et al. [28] scanned and geo-referenced, while the data from 2001-2019 were taken from the FHAAST IDS database compiled by Forest Health Protection, a branch of the Forest Service (https://www.fs.usda.gov/foresthealth/applied-sciences/mapping-reporting/detection-surveys.shtml). Because the 2001-2019 data are vector-based while the 1979-2000 data are rasterized, Dr. Liebhold kindly converted the vector-based data to a rasterized format so that we could more easily compare both types of data to our model projections.

The second data set, for the province of Ontario, is based on data provided by the provincial government (https://geohub.lio.gov.on.ca/documents/lio::forest-insect-damage-event/about) that included the years 1996-2019. Because at least two spongy moth outbreaks occurred in Ontario before 1996 [29], the Ontario data likely underestimate the area of defoliation in that province at least slightly; as fig. 3 shows, however, the existing data cover a reasonably large area. Moreover, we are only using the data to indicate approximately where previous defoliation occurred, and so the Ontario data are sufficient for our purposes. The defoliation map based on the combined USA-Ontario data is then shown in fig. 3.

To use the defoliation data to guide the construction of our model-projection maps, we first had to ensure that the defoliation data were compatible with our model projections. To do this, we re-projected the spongy moth defoliation data on a latitude/longitude scale in the WGS84 coordinate reference system. We then filtered the model-projection locations according to their proximity to defoliation-data locations.

To carry out this filtering, we needed to solve two problems. To begin with, the defoliation data were recorded in 1 km $\times$ 1 km squares, while the model projections were made in 12 km $\times$ 12 km squares, a scale determined by the heavy computing requirements of the dynamically downscaled climate model [12]. The first problem was therefore that we had to identify model-projection squares that were close enough to defoliation squares to justify the inclusion of the model-projection squares in our

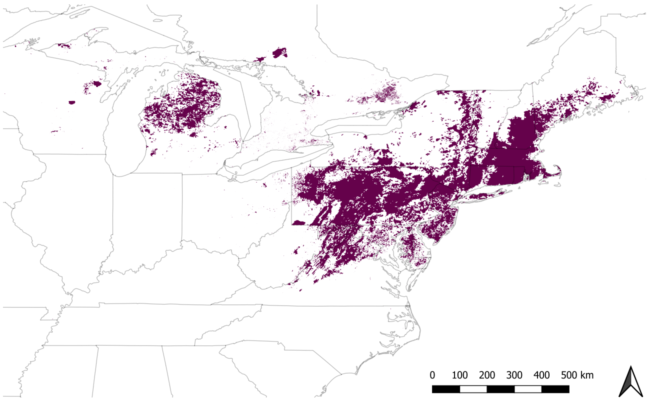

**Fig. 3** Map of spongy moth defoliation in the USA and Ontario, Canada, from 1975 - 2019. Purple shading shows areas that have experienced 1 or more defoliation events. Ontario data are only available beginning in 1996.

maps. Making projections in model-projection squares that were far from defoliation-data squares would have produced over-estimates of the area at risk, whereas making projections only in model-projection squares that are very close to defoliation-data squares would have produced under-estimates of the area at risk. To generate realistic maps, we therefore had to calibrate our filtering process.

The second problem was that the distance encompassed by a degree of longitude depends on the latitude, and so using the Euclidean distance to measure the proximity between a defoliation square and a model square would have resulted in a metric that varied across latitudes, thereby distorting our maps. We therefore instead used a proximity metric that is equal to the sum of the absolute difference in latitude and the absolute difference in longitude between each model square and the nearest defoliation square;

$$G_i = min_j(|\mathrm{Lat}_{M,i} - \mathrm{Lat}_{D,j}| + |\mathrm{Long}_{M,i} - \mathrm{Long}_{D,j}|). \tag{12}$$

Here $\mathrm{Lat}_{M,i}$ is the latitude of the center of model square $i$ ($M$ for "Model"), while $\mathrm{Lat}_{D,j}$ is the latitude of the center of defoliation square $j$ ($D$ for "Defoliation"), while $\mathrm{Long}_{M,i}$ and $\mathrm{Long}_{D,j}$ are the corresponding longitudes of the centers of the two squares. The quantity $G_i$ is thus the total number of degrees between the center of each model square and the center of the nearest defoliation square.

In practice, we selected model squares for which $G_i \leq 0.05$, thereby filtering out model-projection squares that were more than 0.05 total degrees away from any defoliation-data squares. In terms of linear distances, one degree of latitude is equivalent to 111 km, while at the latitudes encompassed by the defoliation data, which range from about 37 to 45 degrees north, one degree of longitude covers between 88 and 75 km. Accordingly, after transformation to a latitude/longitude scale, our 12 km $\times$ 12 km model grid squares encompass approximately 0.11 degrees of latitude and 0.14 to 0.16 degrees of longitude. A value of $G_i \leq 0.05$ is thus equivalent to a distance that is substantially smaller than the width or length of a model square, and so using $G_i \leq 0.05$ ensured that each model-projection square contained at least one defoliation square.

As fig. 4 shows, $G_i \leq 0.05$ does indeed produce maps that visually resemble the defoliation map in fig. 3. Using $G_i \leq 0.02$ in contrast produces maps that visually under-represent the area of previous defoliation (fig. 5), while $G_i \leq 0.2$ produces maps that over-represent the area of previous defoliation (fig. 6). Notably, however, almost all of the under-representation or over-representation is in regions in which previous defoliation has occurred in only a few locations, such as in northern New York State, northern Maine, and the upper peninsula of Michigan. In such regions, a small change in the number of model-projection squares represents a relatively large increase in the area over which we make projections, in turn leading to changes in our maps that are visually obvious. In contrast, for most of the rest of the range of the spongy moth defoliation has occurred in locations that are close together; accordingly, in most of the range a small change in the number of model-projection squares leads to changes that are visually undetectable.

Although changes in $G_i$ thus lead to changes in the visual appearance of our maps, these changes in the visual appearance of the maps do not lead to meaningful changes in the frequency distribution of model outcomes (histograms in figs. 4-6); indeed, the differences in the summary figures in fig. 7 are so small that they are likely due to the necessity of using a finite number of model realizations rather than to differences in the threshold. We therefore conclude that even moderately large changes in $G_i$ have negligible effects on our overall results, and thus that our model-projection maps provide a reasonable approximation to the defoliation-data map.

### 4 Comparison of Weather in our Study Plots to Weather Across the Range of the Spongy Moth

As we mentioned in the main text, the parameters of our eco-climate model were estimated from data that we collected in the lower peninsula of Michigan during the spongy moth larval seasons in 2010, 2011 and 2012 [1]. Because the study sites were located along a 300 km-long north-south transect, our weather data include substantial variation in each weather variable. As fig. 8 shows, this variation encompasses most of the variation in the same weather variables for the same period across the range of the spongy moth in eastern North America; moreover, the mean of each variable is similar to the corresponding mean calculated across the range of the spongy moth. This similarity suggests that model projections based on parameters estimated in Michigan are likely to hold for other parts of the range of the spongy moth.

### 5 Alternative Scenarios

Here we consider four alternative scenarios. In the first alternative scenario, we include nonlinear effects of increasing temperatures on spongy moth larval growth and survival; in the second, we vary the initial density of resting spores; in the third, we vary the climate-change scenario; in the fourth, we make projections for the middle of the century instead of the end of the century. For the first two scenarios we use the same climate-change maps as in the main text, and so to avoid redundancy in those cases we show summaries of model outcomes but not maps.

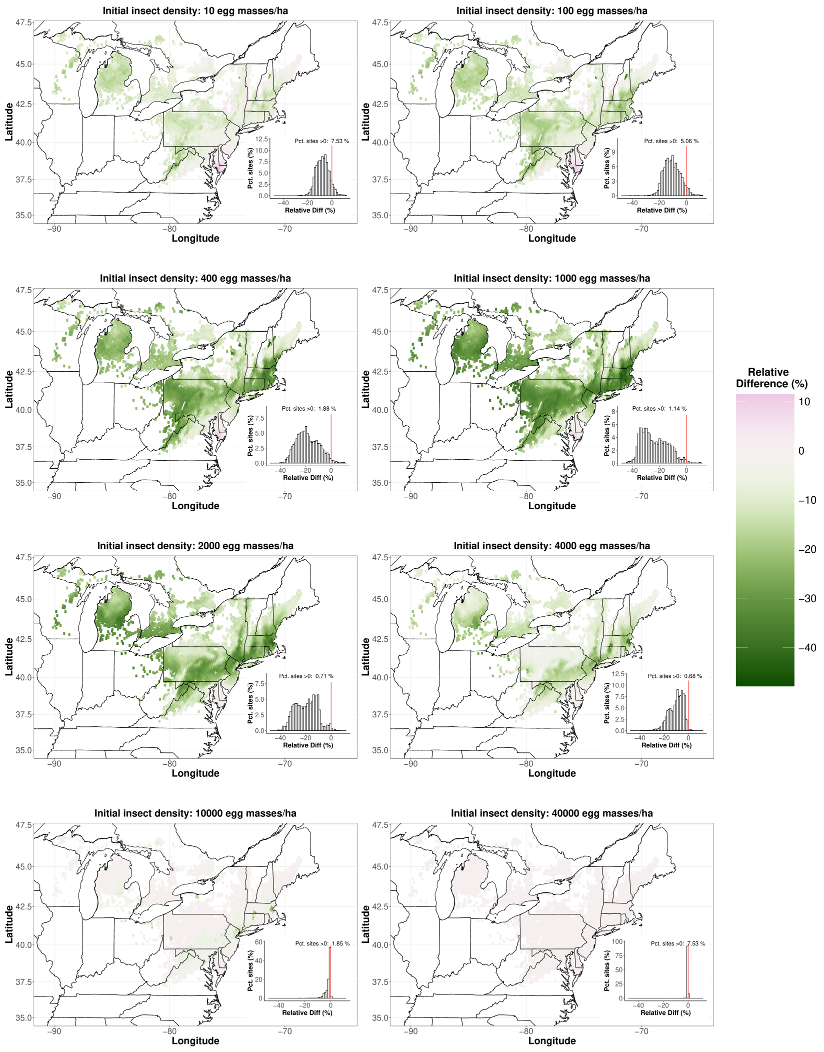

**Fig. 4** A copy of fig. 2 in the main text, showing changes in the percent infected with *E. maimaiga*. In this case, the threshold value of our distance metric $G_i \leq 0.05$, with $G_i$ calculated as in equation (12). These maps are presented here for comparison to the maps in figs. 5 and 6, which respectively under-represent and over-represent the areas of previous defoliation.

#### 5.1 Reductions in Larval Growth Rates and Survival Due to Heat Stress

As we explained in the main text, to ensure that our model projections are statistically robust we used field data to select the best model from a group of 15 competing models [1]. Best practice in model selection dictates the avoidance of models for which there is little *a priori* support from the data [30], and so we considered only models in

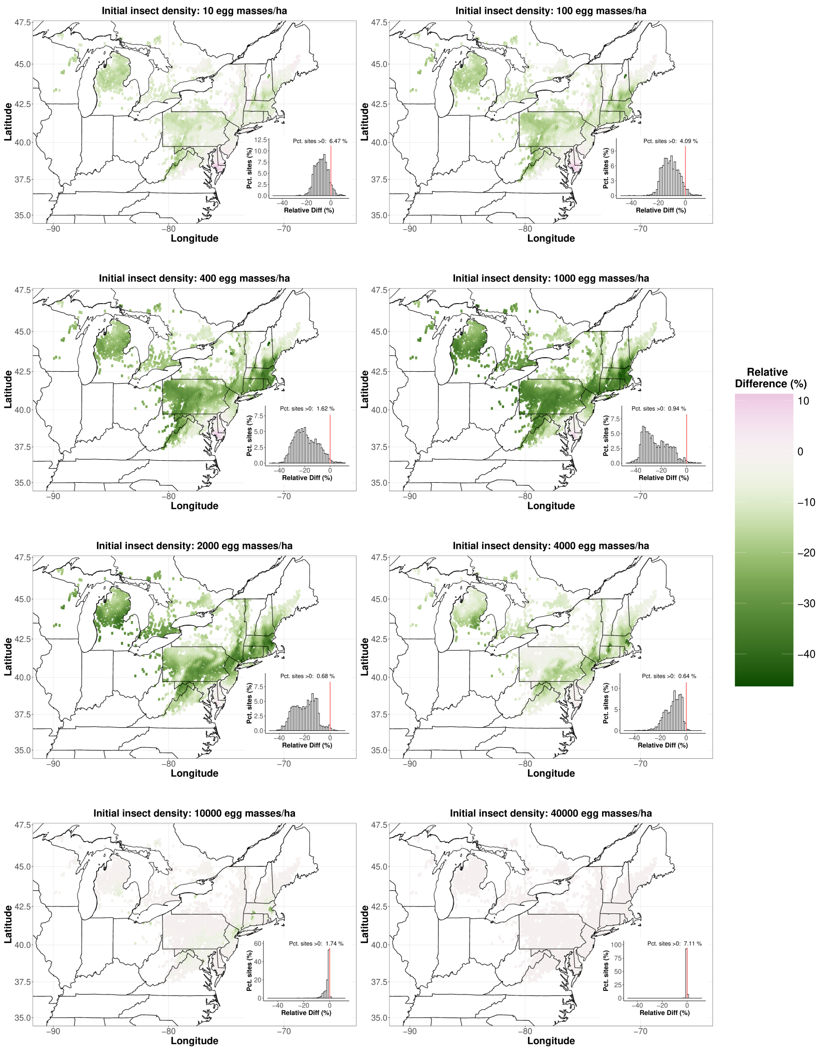

**Fig. 5** Maps of the change in the percent infected with *E. maimaiga* constructed using a value of the threshold $G_i \leq 0.02$. Comparison with fig. 4 shows that this value of $G_i$ leads to a visual under-representation of the area of defoliation shown in fig. 3.

which larval size increases linearly with temperature. As we mentioned earlier, however, laboratory experiments at constant temperatures have recorded either declines or at least an end to increases in larval growth at roughly 28° C [18], suggesting that the assumption of linear growth is incorrect. It is nevertheless also true that when higher temperatures during the day are accompanied by lower temperatures at night, then growth rates do not begin to decline until the daytime temperature is above 36 to 38° C [19]. In the latter case, the temperature at which the decline occurs differs depending on the length of time for which daytime temperatures are high; if the temperature

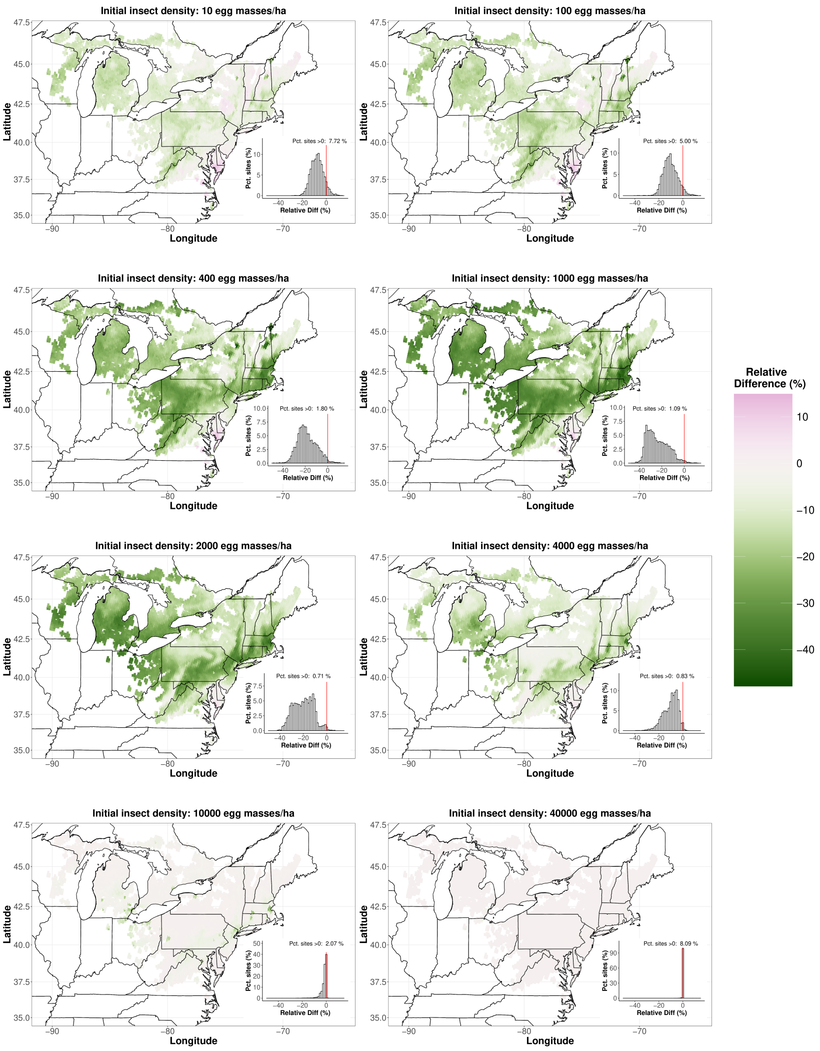

**Fig. 6** Maps of the change in the percent infected with *E. maimaiga* constructed using a threshold value of the distance metric of $G_i \leq 0.2$. Comparison with fig. 4 shows that this value of $G_i$ leads to a visual over-representation of the area of defoliation shown in fig. 3.

during the daytime is high for 7 days, then the decline starts at a daytime temperature of 36° C, but if the temperature during the daytime is high for 2 days, then the decline does not start until the daytime temperature reaches 38° C [19]. Constant and variable temperatures thus give very different results in the laboratory, and even modest changes in variable-temperature experiments lead to detectable differences in model outcomes. Meanwhile, even laboratory experiments with fluctuating temperatures likely provide only a rough approximation to field temperatures. In the main

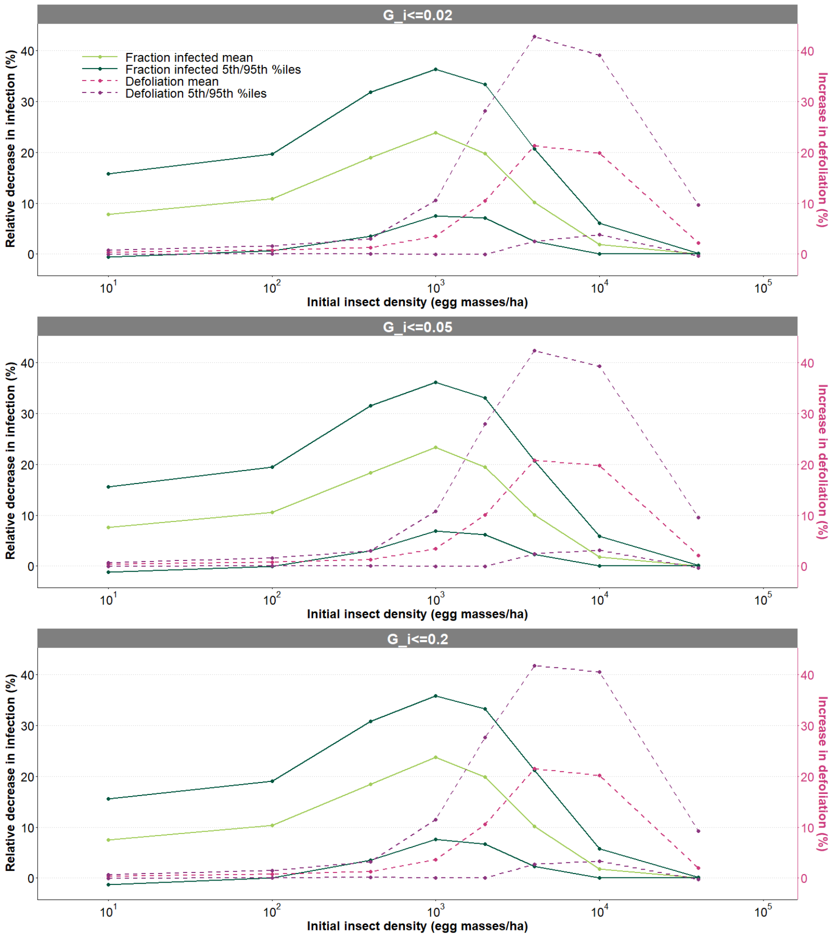

**Fig. 7** Summary plots for different threshold values of our distance metric $G_i$. The middle plot is identical to fig. 3 in the main text except that here, and for subsequent summary plots, we remove the histograms, we put the change in the defoliation rate on the right vertical axis, and we change the range of the vertical axes. As in the main text, we multiply the change in the fraction infected by $-1$ so that changes in the fraction infected and changes in defoliation have the same sign. In the top plot we use $G_i \leq 0.02$ (corresponding to fig. 5) and in the bottom plot we use $G_i \leq 0.2$ (corresponding to fig. 6). The very small differences between plots make clear that moderate changes in $G_i$ have essentially no effect on our results.

text, we therefore present results only for a model fit to field data; as we described, this model assumes that larval growth increases linearly with temperature.

Another reason why we focus on the linear growth model is because it is conservative, in the following sense. In our model and in nature, the *E. maimaiga* transmission rate is higher when larvae are larger; the assumption that larval size increases linearly with temperature thus means that, all else equal, increased temperatures due to climate change will result in increases in the *E. maimaiga* infection rate, thereby lessening the negative effects of climate change that are the focus of our work. The

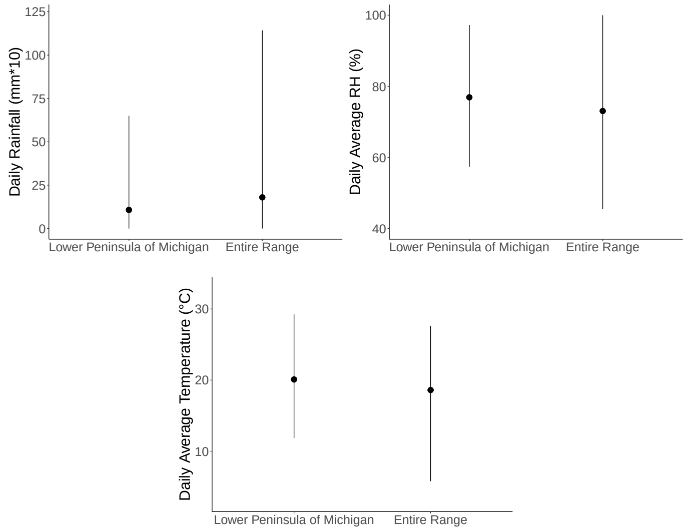

**Fig. 8** Comparison of weather data from our study plots to weather data from the National Center for Environmental Information for the range of the spongy moth. Points indicate means and bars indicate upper 95th and lower 5th percentiles of the distribution of each variable. The similarity in means and in upper 95th/lower 5th percentiles makes clear that the weather conditions during our field study in Michigan were similar to the weather conditions for the same period across the range of the spongy moth.

climate-change driven decline in the *E. maimaiga* infection rate in our model thus occurs in spite of our assumption that larval size increases linearly with increasing temperature.

High temperatures, however, are likely to reduce not just larval growth rates, but also larval survival, again because of the heat stress associated with high temperatures [31]. Although our model also does not include increases in non-disease larval mortality with increased temperatures, such increases could in principle weaken our conclusions. This is because deaths due to high temperatures reduce the spongy moth density, thereby lowering the *E. maimaiga* infection rate and increasing the spongy moth defoliation rate. Allowing for deaths due to heat stress might therefore reduce the effects of climate change, and so eliminating the assumption that larval survival is unaffected by increasing temperatures could weaken our conclusions and may thus be anti-conservative.

To test for this possibility, we expanded our model to allow for increased non-disease mortality at high temperatures; for completeness, we also allowed for reduced larval growth rates at high temperatures. Out of concern that the laboratory data were unreliable, we also considered a second scenario that allowed for more severe reductions in non-disease mortality than were seen in the laboratory data.

In these models, we assumed that at temperatures above 28° C the larval growth rate was equivalent to its value at 28° C. This assumption roughly splits the difference between the Logan et al. [18] study of laboratory growth at constant temperatures, in which growth rates decline slightly at temperatures above 28° C, and the Banahene et al. [19] study of laboratory growth at varying temperatures, in which growth rates instead did not begin to flatten out until temperatures reached either 36° C or 38° C. We repeat that assuming that growth sharply declines with increased temperature would be anti-conservative, so allowing for a leveling off of the growth rate at high temperatures is a more conservative assumption.

To also incorporate increases in larval non-disease mortality at high temperatures, we added temperature-driven non-disease mortality to our model, in the same way that we had earlier added temperature-driven conidia mortality. (As we mentioned earlier in this document, our parameter values implicitly incorporate the effects of the non-disease mortality that occurs between when larvae hatch and when they finish dispersing off their egg masses onto foliage.) We thus added an additional temperature-dependent mortality function to the susceptible-host and exposed-host equations in the model. To construct this mortality function, we began with a model that Thompson et al. [31] created to describe larval mortality at constant temperatures in the laboratory. As we described, however, the data from Banahene et al. [19] instead quantify mortality at varying temperatures and are thus closer to the situation in the field. We therefore used the Thompson et al. model to describe larval survival as a function of temperature, but we fit the model's parameters to the Banahene et al. data.

In the Thompson et al. model, when the temperature is $T_\tau$ on day $\tau$, the probability of larval survival $P_{S,\tau}$ is:

$$P_{S,\tau} = \frac{1}{1 + exp\big(-f(T_\tau)\big)}, \tag{13}$$

Here $S$ stands for survival, and the function $f(T_\tau)$ is;

$$f(T_\tau) = \beta_0 + \beta_1 T_\tau + \beta_2 T_\tau^2. \tag{14}$$

For Thompson et al.'s purposes, this function is useful because $f(T_\tau)$ is the logit-transformed mortality, and so the combination of equations (13) and (14) allowed Thompson et al. to analyze their data using standard logistic regression software. Because our experience has been that such software does not always converge on the best parameter set, we instead used the nonlinear fitting routine `optim()` in the `R` programming language. Like Thompson et al., however, we used a binomial likelihood function, and we used maximum likelihood to fit the model to the data. To ensure that our fitting routine had converged, we repeated it using $10^5$ random restarts, such that the initial parameters for each restart were drawn randomly from an area of parameter space in the vicinity of a parameter set that we generated in an initial fitting attempt. The top 10% of these restarts gave effectively identical parameters, confirming the robustness of our fitting routine. Reassuringly, as fig. 9 shows, the model gives an excellent fit to the data; moreover, as we will show, changing the parameters of this model had almost no effect on our results in any case.

To consider an even more conservative version of the model, we produced an alternative set of parameters by reducing survival at each temperature in the Banahene et al. study, and then re-fitting the model to the data. To preserve the shape of the survival function, we reduced survival in such a way that the shape of the survival

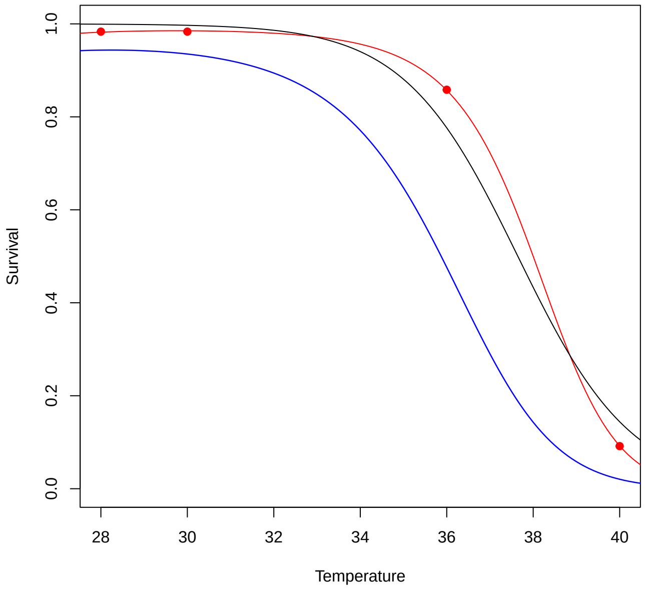

**Fig. 9** Models fit to data on survival versus temperature from Banahene et al. [19]. The red circles represent the data, the red line shows the best-fit version of equations (13)-(14), and the black line shows a best-fit version of equations (13)-(14) for which $\beta_2 = 0$. The AIC score for the $\beta_2 = 0$ model is higher by 18.6 points, confirming that the $\beta_2 \neq 0$ model fits the data much better than the $\beta_2 = 0$ model. For the red line, the maximum likelihood estimates of the parameters, with bootstrapped 95% confidence intervals in parentheses, are $\beta_0 = -50.631$ $(-75.508, -18.090)$, $\beta_1 = 3.685$ (5.156, 1.901), $\beta_2 = -0.0619$, $(-0.0834, -0.0370)$. The blue line shows an alternative version of the red line, for which we arbitrarily changed the model parameters to $\beta_0 = -35.762$, $\beta_1 = 2.732$, $\beta_2 = -0.0484$. This latter model thus imposes lower survival while having roughly the same shape as the red line. Because the data show no effects of heat stress on survival at temperatures less than 28° C, all of our models assume that there is no heat-stress induced mortality at temperatures below 28° C.

function was roughly the same but survival fell more rapidly with increasing temperature. The higher mortality in the model based on these synthetic data provided an additional and more severe test of the effects of heat-stress-driven mortality on our results.

To produce a mortality function for our model, we translated the survival function $P_{S,\tau}$ in equation (13) into an instantaneous mortality rate by assuming that mortality on day $\tau$ is the result of a constant mortality rate on that day, according to:

$$\mu_{H,\tau} = -\frac{1}{20} \log\big(P_{S,\tau}\big). \tag{15}$$

Here $H$ stands for heat-stress, so that the heat-stress-induced death rate $\mu_{H,\tau}$ is distinct from the pathogen-induced death-rate $\mu_{C,\tau}$ in the full model. Although the time that it took larvae to develop varied with temperature in the Banahene et al. study

[19], we assumed a constant development time of 20 days, which is roughly the lowest development time observed in that study. Given that our goal is to test whether non-disease mortality affects our conclusions, this relatively short development time leads to a conservatively high mortality rate.

We then generated a new set of model projections using mortality functions based on both the original Banahene et al. data and our synthetic data. These projections show that allowing for lower larval growth and higher non-disease larval mortality at higher temperatures has essentially no effect on our model's projections (fig. 10). We therefore conclude that the ecological effects of climate change on the spongy moth-*E. maimaiga* interaction are likely to play a more important role in determining spongy moth defoliation than the physiological effects of temperature on the spongy moth.

There are two reasons why heat-stress effects on larval growth and survival have no effect on our results. First, temperatures that are high enough to cause strong effects on growth and survival do not occur until late in the larval season. This is important because the trajectory of the *E. maimaiga* epizootic is largely determined by events over the first half of the larval season [1], and so reductions in larval growth and survival late in the season have little effect on the epizootic. Second, although our climate-change model projects that temperatures will rise across almost the entire range of the spongy moth, temperatures will generally increase by less than a degree; as fig. 9 makes clear, increases of this amount are likely to produce only very modest declines in larval growth and survival.

#### 5.2 High and Low Resting Spore Densities

Little is known about the determinants of variation in resting spore densities [13], and so we assumed that resting spore densities are constant over time and space. As we described earlier, the resting spore densities that we used in our models then came from our previous work, in which we studied populations in the early stages of collapse [1], whereas resting spore densities tend to be higher in spongy moth populations that have already collapsed. Because here we are instead concerned with whether *E. maimaiga* can begin the process of collapse, our previous estimates are likely to provide the most useful projections of future infection rates. For the eco-climate model projections in the main text, we used a resting spore density of $5.71 \times 10^{-3}$ m$^{-2}$, which is the second highest of our seven estimates. Because our overall argument is that climate change is likely to reduce *E. maimaiga* infection rates, and because as we will show high resting spore densities ameliorate the effects of climate change, using a higher value is conservative. The variation among our estimates was not particularly high, however, with a standard error of $2.38 \times 10^{-3}$ m$^{-2}$, so our default value is within one standard error of the mean value of $4.75 \times 10^{-3}$.

To show the effects of resting-spore variation, here we show cases in which the resting-spore density is instead either the second-lowest of our estimated values, $6.91 \times 10^{-4}$ m$^{-2}$ (the lowest value was an extreme outlier), or the highest of our estimated values, $1.83 \times 10^{-2}$ m$^{-2}$. At more than 2 standard errors higher than the mean, the highest value is quite extreme, which is why we used the second highest value in the main text.

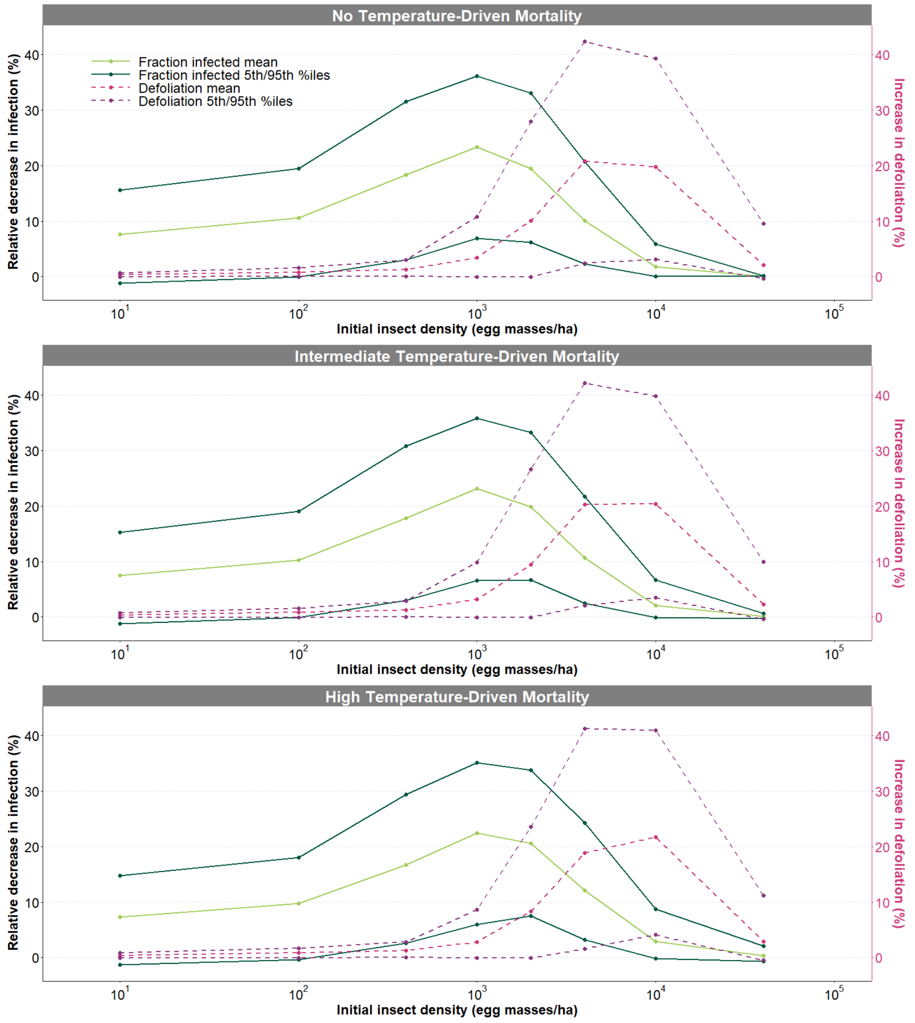

**Fig. 10** Effects of heat-stress-induced reductions in larval growth and survival on our model's projections. The top plot is based on fig. 3 in the main text, for which we assumed that there is no non-disease mortality. The middle plot instead includes heat-stress-driven mortality, based on the red curve in fig. 9, while the bottom plot includes heat-stress-driven mortality based on the blue curve in fig. 9. The similarity of the three plots demonstrates that adding reduced growth and increased non-disease mortality to our model has no effects on the model's projections.

As the top panel of fig. 11 shows, the highest resting spore density leads to moderately less severe reductions in the infection rate than the second-highest density (middle panel of fig. 11). Meanwhile, the bottom panel of fig. 11 shows that the second-lowest resting-spore density in contrast leads to much more severe reductions in the infection rate than the second-highest density. Fig. 11 further shows that the increases in the defoliation rate roughly match the reductions in the infection rate. Lower resting-spore densities thus lead to even bigger negative effects of climate change, which is why our use of the second-highest density in the main text is conservative.

The effects of climate change on the infection rate are weaker at higher resting spore densities because in both the model and in nature resting spores are responsible for most or all *E. maimaiga* infections at the beginning of epizootics: high densities of resting spores therefore lead to more severe epizootics. Meanwhile, because of the non-linearities inherent in the conidia-transmission process, climate change has a stronger effect on conidia transmission than on resting spore transmission, and so higher resting spore densities can partially compensate for the negative effects of climate change. Lower resting spore densities in contrast exacerbate the negative effects of climate change. This is because when resting spore densities are low conidia play a bigger role in determining epizootic severity, and the strong effect of climate change on conidia then leads to sharp reductions in the infection rate.

More generally, fig. 11 makes clear that reductions in the *E. maimaiga* infection rate and increases in the defoliation rate are still severe when the resting spore density is high. We thus conclude that climate change would still have meaningfully negative impacts on *E. maimaiga* epizootics even if resting spore densities in nature are consistently higher than the resting spore densities that we used in our model. If resting spore densities are instead consistently lower than the resting spore densities that we used in our model, then the impacts of climate change on the *E. maimaiga* infection rate would be even more severely negative than shown in the main text. We therefore argue that our conclusions are robust to variation in resting spore density.

A final point is that the importance of spongy moth density for the conidia transmission process means that the effects of climate change are more strongly altered by variation in spongy moth densities than by variation in resting-spore densities [1]. In the main text we therefore focus on variation in spongy moth densities.

#### 5.3 RCP 4.5

Here we show results for the RCP 4.5 climate-change scenario, a more optimistic alternative to the RCP 8.5 scenario that we used in the main text. As a reminder of the RCP 8.5 case, in fig. 12 we first show the RCP 8.5 climate-change map from the main text, while in fig. 13 we show the corresponding RCP 4.5 map. The projected changes in the two cases are similar, in that rainfall rises in more than 30% of locations in both cases, while relative humidity falls over most of the landscape and temperature increases over most of the landscape in both cases. The general trends of climate change are thus roughly similar for the two cases.

Notably, however, the frequency distributions of outcomes across locations for the two cases have quite different shapes, in a way that matters. Specifically, in the RCP 4.5 case the means and the upper 95th and lower 5th percentiles of the changes in rainfall and relative humidity indicate more severe reductions than in the RCP 8.5 case, and in the RCP 4.5 case the fraction of locations that show increases in temperature is considerably higher than in the RCP 8.5 case. It is nevertheless true that in the RCP 8.5 case the upper 95th percentile on the increase in temperature is considerably higher, with a cluster of locations showing sharp increases in temperature, and with a high frequency of sharp reductions in relative humidity. Because of these latter

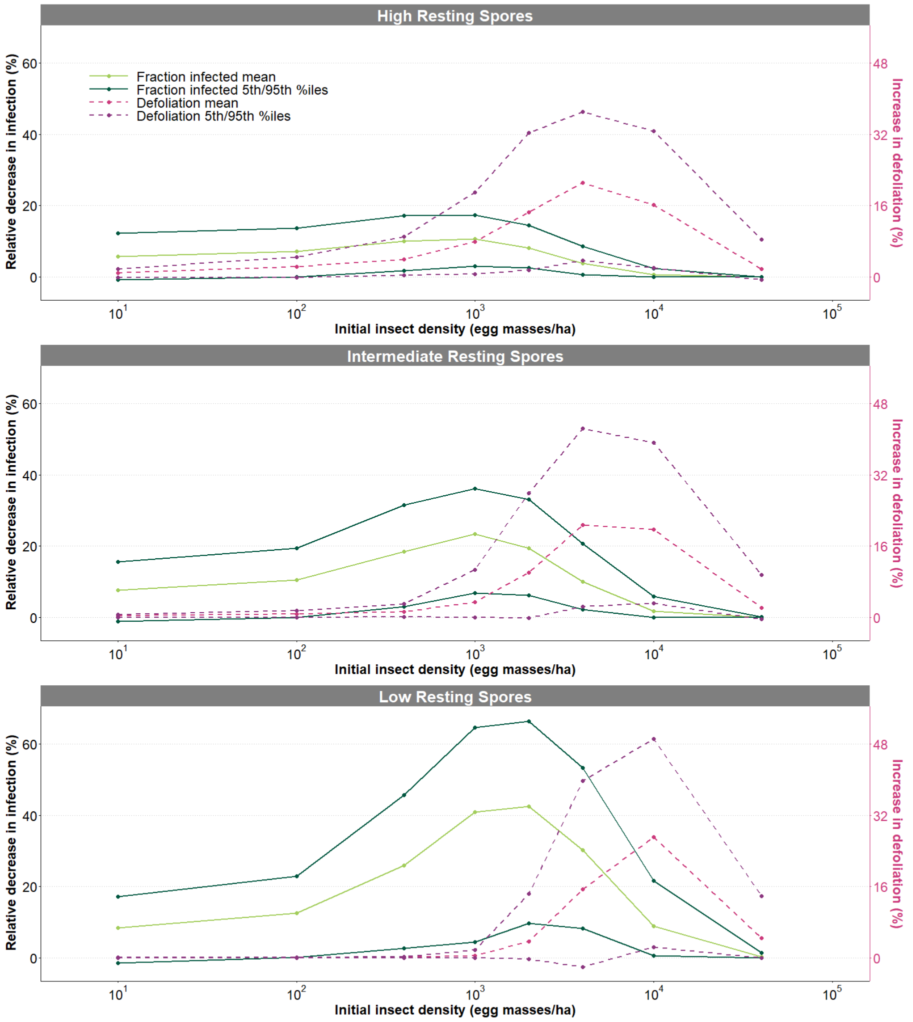

**Fig. 11** Summary plots of the effects of resting spore densities on our model's projections. Here we use a similar format to the plots in fig. 21, so that the middle plot is identical to fig. 3 in the main text. In the top plot we increased the resting spore density to $1.83 \times 10^{-2}$ m$^{-2}$ relative to the value of $5.71 \times 10^{-3}$ m$^{-2}$ that we used in the main text, while in the bottom plot we reduced the resting spore density to $6.91 \times 10^{-4}$ m$^{-2}$. A lower resting spore density than the value that we used in the main text thus leads to more severe effects of climate change, while a higher resting spore density instead leads to moderately less severe effects of climate change.

complications, the RCP 4.5 case generally leads to moderately less severe change in the fraction infected and in defoliation than the RCP 8.5 case (figs. 14-16). At the lowest spongy moth densities, however, the extent of the reduction in the infection rate under the RCP 4.5 scenario is slightly stronger than under the RCP 8.5 scenario, which is important because of the importance of *E. maimaiga* epizootics at low spongy moth densities. The overall effect, however, is that the differences in the two cases are

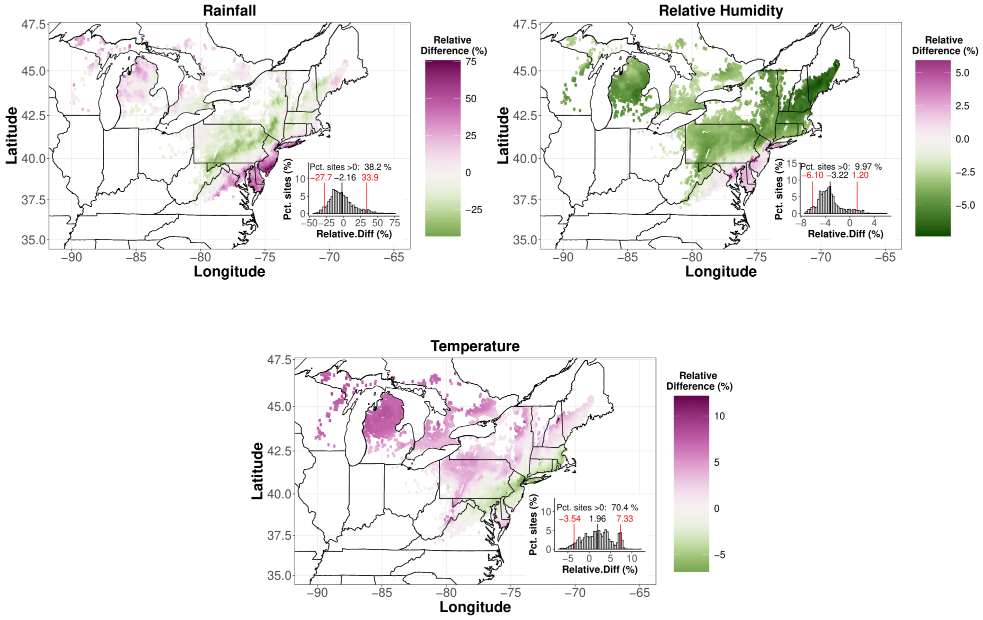

**Fig. 12** Projections of relative changes in rainfall, relative humidity, and temperature for our climate-change model under the RCP 8.5 scenario at the end of the century. This is a copy of fig. 1 from the main text that is placed here for comparison with fig. 13 and fig. 17 below, which differ in showing projections under the RCP 4.5 scenario and for the middle of the century, respectively.

not that large, and so we conclude that our results are robust to changes in the $CO_2$ concentration scenario.

#### 5.4 Mid-century Projections

Here we present projections for the mid-century, specifically for the period 2045-2054, as opposed to the end-of-century projections in the main text, which instead covered the period 2085-2094. In fig. 17 we show mid-century changes for the weather, as projected by our climate-change model; because here we again use the RCP 8.5 scenario, this figure can be compared to fig. 12, which shows end-of-century projections under the RCP 8.5 scenario. The main difference is that the mid-century projections show greater increases in rainfall at the northwest edge of the insect's range, and smaller declines in relative humidity across the landscape.

Because the spongy moth-*E. maimaiga* interaction is quite sensitive to weather, however, the smaller declines in relative humidity at mid-century lead to reductions in the infection rate that are almost as severe as the reductions in the infection rate at the end of the century, and the reductions again occur almost everywhere (fig. 18). The frequency of areas that will experience increases is moderately higher than at the end of the century, but the increases are generally small and are restricted to a small fraction of the landscape. Meanwhile, in locations where declines in the infection rate occur,

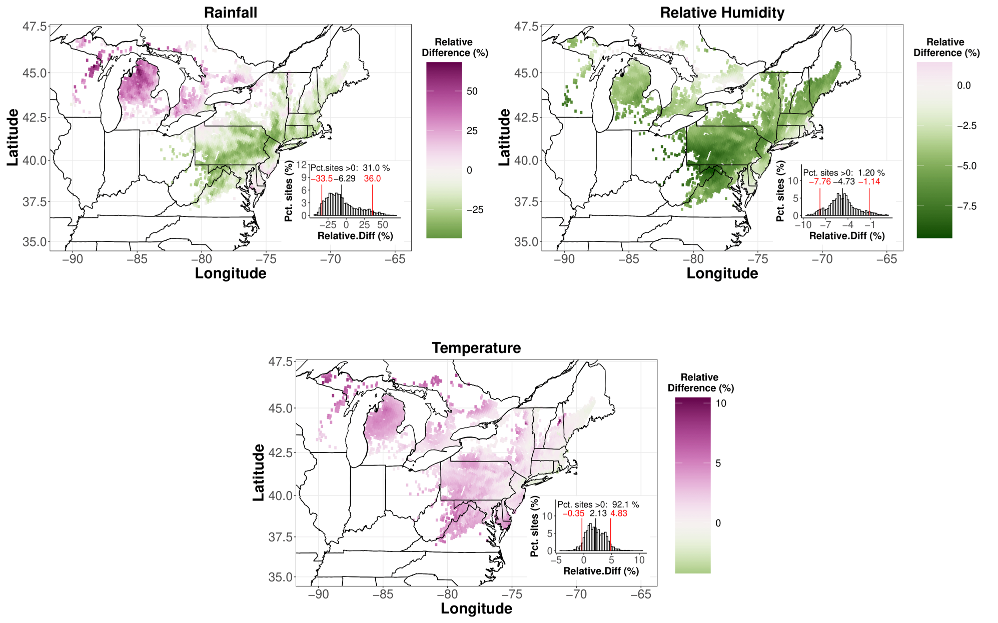

**Fig. 13** Projections of relative changes in rainfall, relative humidity, and temperature for our climate-change model under the RCP 4.5 scenario, as opposed to the RCP 8.5 scenario that we used in fig. 12 above. As the inset histograms show, this scenario differs in complex ways from the RCP 8.5 scenario.

the declines in the infection rate and the corresponding increases in the defoliation rate (fig. 19) are almost as severe as the corresponding declines and increases that are projected to occur by the end of the century. Compared to the end-of-century projections, the mid-century projections thus show a modest rise in the frequency of increases in the fraction infected, while nevertheless retaining a high frequency of severe declines, with corresponding changes in defoliation (fig. 20).

### 6 Effects of Stochasticity on the Model Projections

In the main text, we consider only average model projections; to instead allow for the effects of stochasticity, here we consider variation in the model outcomes. Although the weather conditions are the same in each model realization, the realizations nevertheless differ because of the stochastic terms in the models. Stochasticity in the models is thus due to inherent stochasticity rather than to variation in weather conditions.

As we have described, we had already generated 100 realizations for each historical year and each future year, and we had 10 years of historical and future scenarios, respectively. We thus had a total of 1000 values for the historical scenarios and 1000 values for the future scenarios. We then used the R function `outer` to calculate the pairwise differences in all $1000 \times 1000 = 10^6$ combinations: using this function made the calculation much faster than if we had instead used a series of `for` loops. As in the main text, we calculated relative percent changes in infection rates, and untransformed

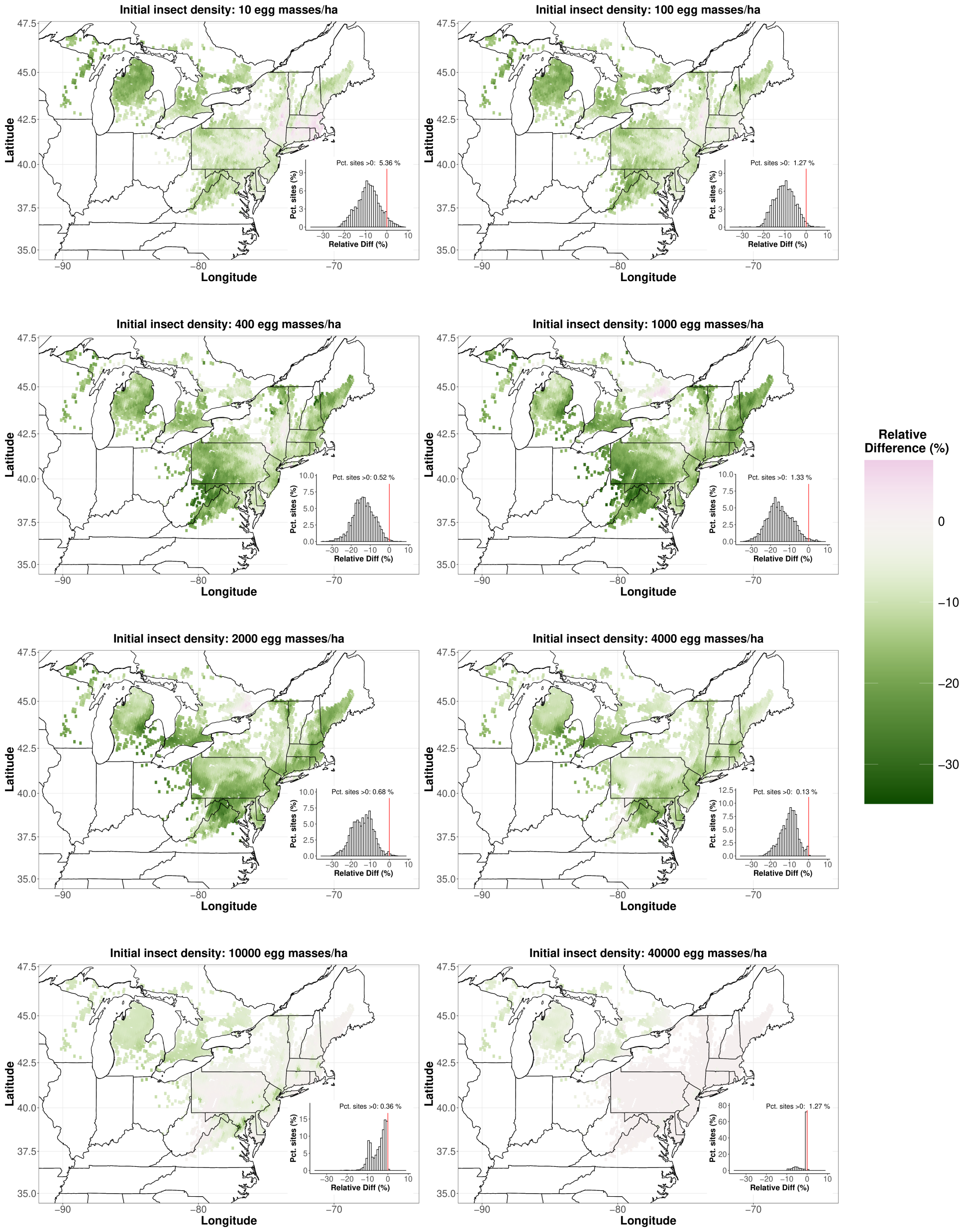

**Fig. 14** Projections of the eco-climate model as in fig. 2 in the main text, except that here we use the RCP 4.5 scenario.

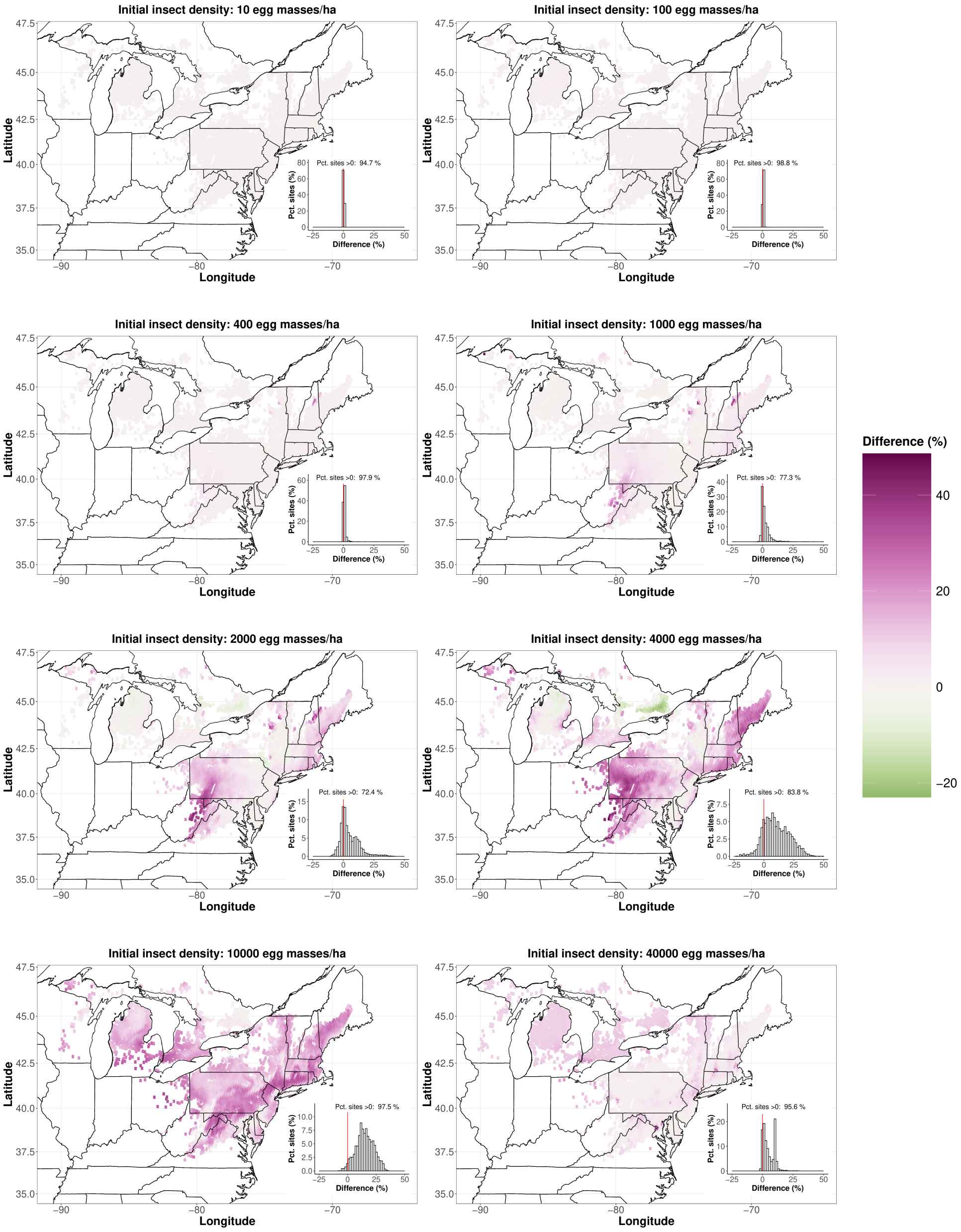

**Fig. 15** Projections of defoliation as in fig. 4 in the main text, except that here we use the RCP 4.5 scenario.

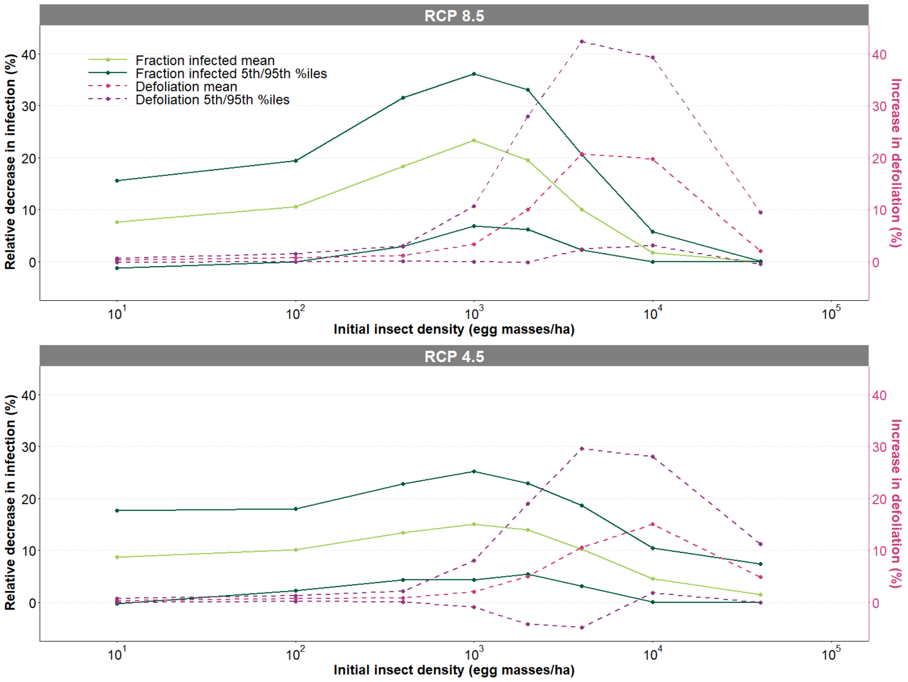

**Fig. 16** Summary of model projections under the RCP 8.5 and 4.5 scenarios. As in earlier figures, the upper plot is identical to fig. 3 in the main text, which used the RCP 8.5 scenario. In the lower plot we instead use the RCP 4.5 scenario, based on values from fig. 14 and fig. 15. The RCP 4.5 case thus leads to slightly less dramatic changes in the fraction infected and in defoliation compared to the RCP 8.5 case, but the differences are modest.

percent changes in defoliation rates. We then calculated the upper 75th and lower 25th percentiles of each statistic across its $10^6$ values at each geographical location. Here the upper percentiles indicate more severe effects of climate change and the lower percentiles indicate less severe effects of climate change.

As in the main text, we summarize the projections in terms of the 5th and 95th percentiles of the distribution of outcomes across geographic locations. Meanwhile, as we described, the outcome at a given location consists of the projections at either the 75th or 25th percentile of outcomes calculated across stochastic model realizations at that location. In what follows, it is therefore important to keep in mind the distinction between the percentiles calculated across locations and the percentiles calculated across realizations.

As the lower panel in fig. 21 shows, the model's projections are moderately less severe at the lower 25th percentile of projections than at the means (middle plot in fig. 21). At the 25th percentiles, climate change leads to increases in the infection rate at almost 50% of sites; over the rest of the landscape, however, infection rates are again reduced, and at intermediate spongy moth densities the extent of the reductions is often quite severe. Sheer luck may thus lead to mild increases in the infection rate about 25% of the time across about 50% of the landscape; over the rest of the landscape, however, not even sheer luck will prevent reductions.

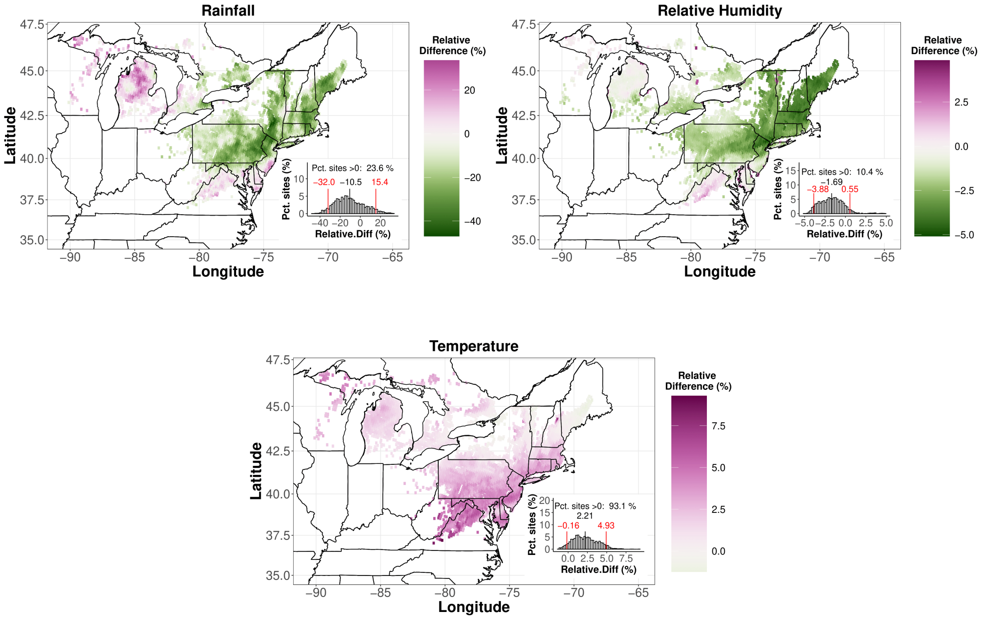

**Fig. 17** Mid-century projections of changes in weather, as opposed to the end-of-century projections in the main text. Comparison to fig. 12 shows that mid-century changes will be moderately less severe than end-of-century changes.

In contrast, as the top panel in fig. 21 shows, at the upper 75th percentiles the model instead shows sharper reductions in infection and sharper increases in defoliation than occur at the means. Sheer luck will thus also lead to much more severely negative outcomes.

The results at the 25th percentiles are thus moderately less severe than the results at the means, whereas the results at the 75th percentile are substantially more severe than the results at the means. This is the basis of our argument in the main text that outcomes that are more severe than the mean are highly likely, whereas outcomes that are less severe than the mean are unlikely. The average projections in the main text are thus optimistic.

### 7 Comparison of Recent and Historical Defoliation Maps

In fig. 22, we compare defoliation in three distinct periods relative to the initial introduction of *E. maimaiga* into North America. Although the initial introduction occurred in 1989, *E. maimaiga* releases continued up to 1996, by which time the pathogen had occupied essentially the entire area of the USA in which significant defoliation has been recorded [32]. Meanwhile, 1996 is the first year for which detailed data are publicly available for Ontario, Canada.

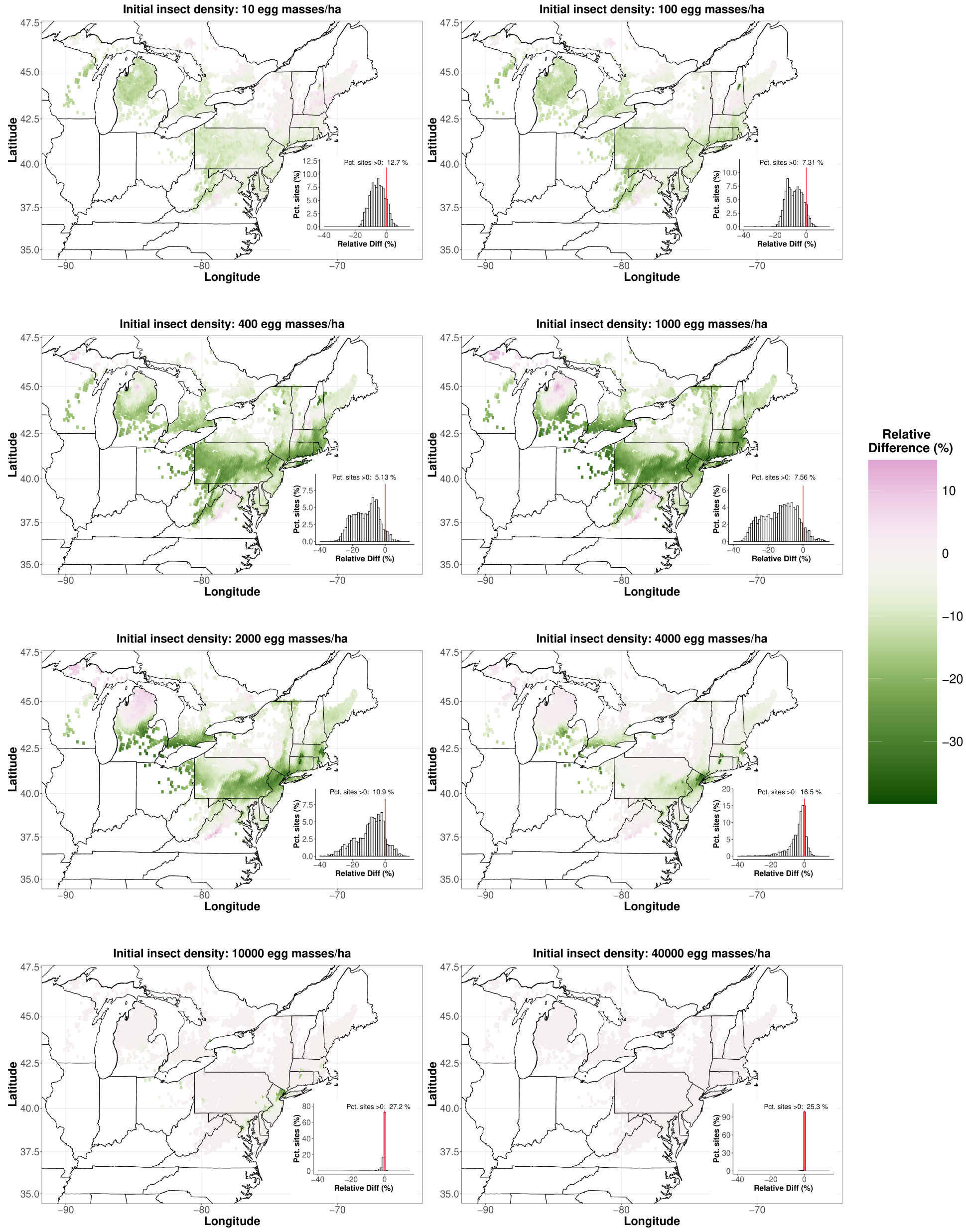

**Fig. 18** Mid-century projections of changes in infection rates.

Here we therefore show defoliation maps for three *E. maimaiga* eras; first, the pre- and early-*E. maimaiga* era, from 1975-1995; second, the era in which *E. maimaiga* kept defoliation at low levels, from 1996-2014; and third, what may be the beginning of an era in which *E. maimaiga* is no longer able to keep defoliation at low levels, from 2015

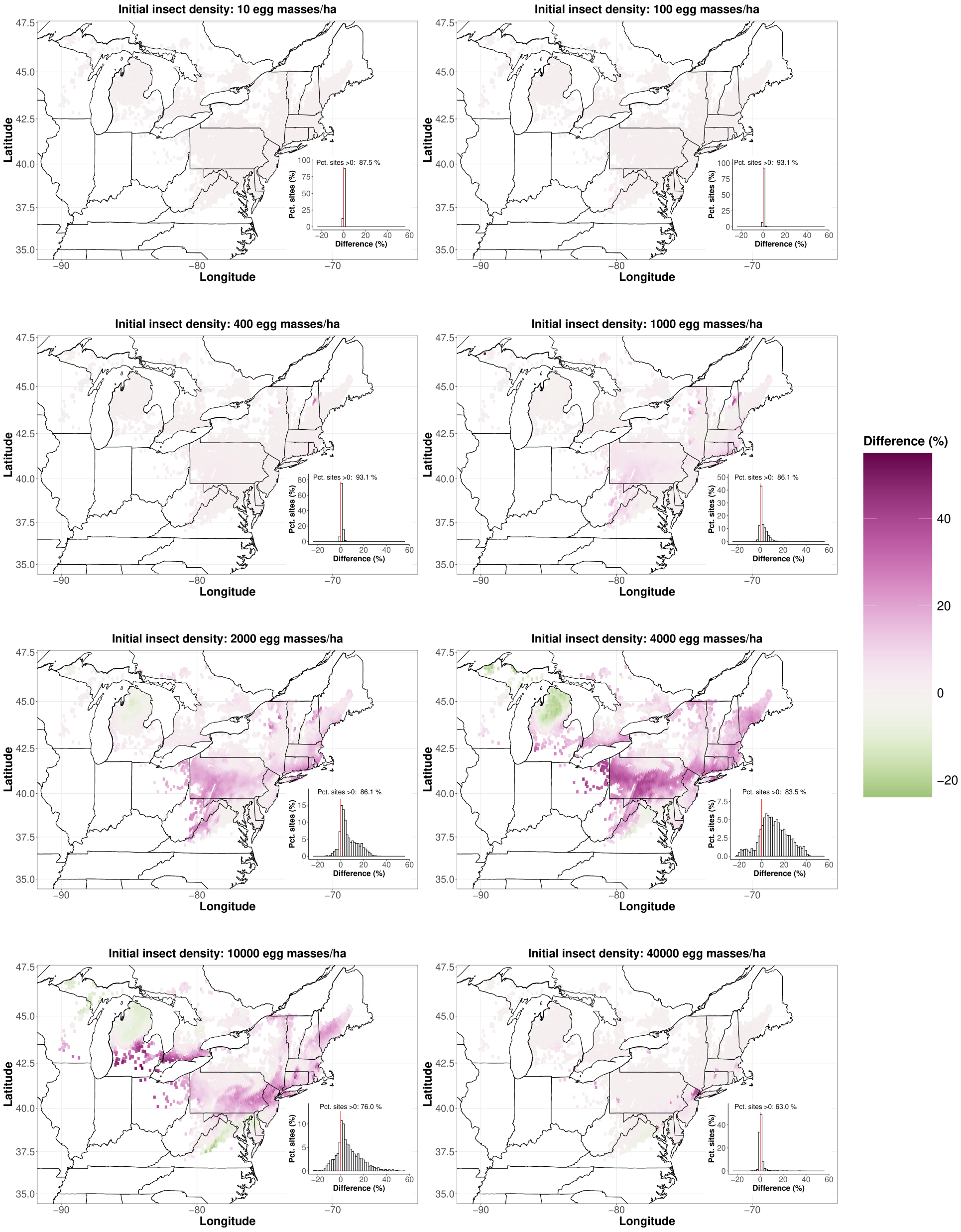

**Fig. 19** Mid-century projections of defoliation.

to the present. We used 2015 as the beginning of the latter period because, as the maps
765 indicate, in 2015-2018 southern New England experienced its first severe outbreak since the introduction of *E. maimaiga* [33], while in 2020-2021 Ontario similarly experienced its first severe outbreak since the introduction of *E. maimaiga* [29]. The 2015-2021 map

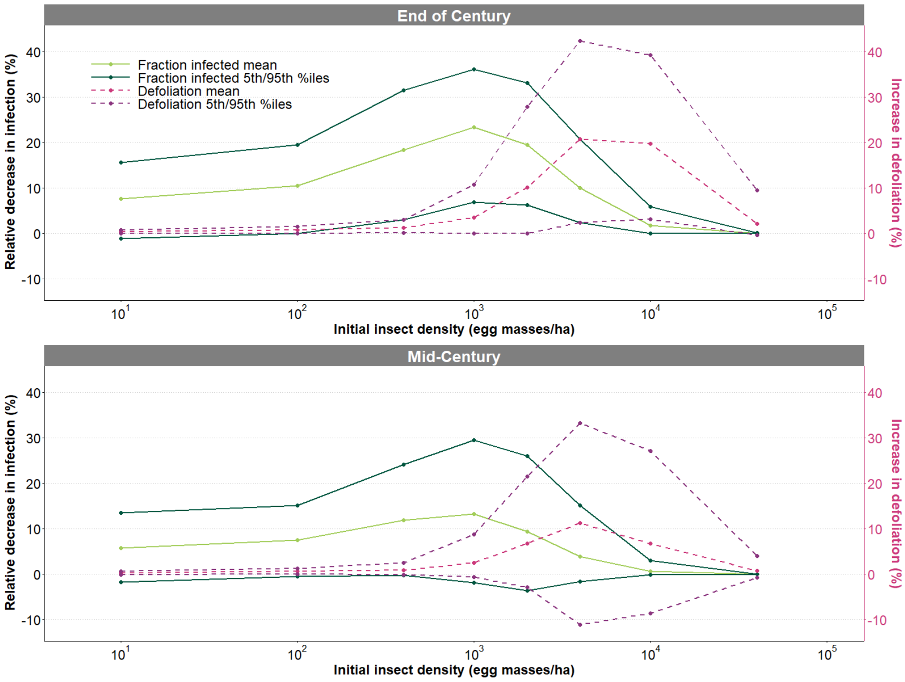

**Fig. 20** Summary of model projections at the end and middle of the 21st century. The upper plot shows the end-of-century projections from fig. 3 in the main text, as in previous summary figures. In the lower plot we show the mid-century scenario, based on the values in fig. 18 and fig. 19. The mid-century projections thus show only modestly less severe reductions in the fraction infected and in defoliation.

also shows the beginnings of an outbreak in northern New England that continued into 2022, but at this writing mapping data are not yet publicly available for 2022.

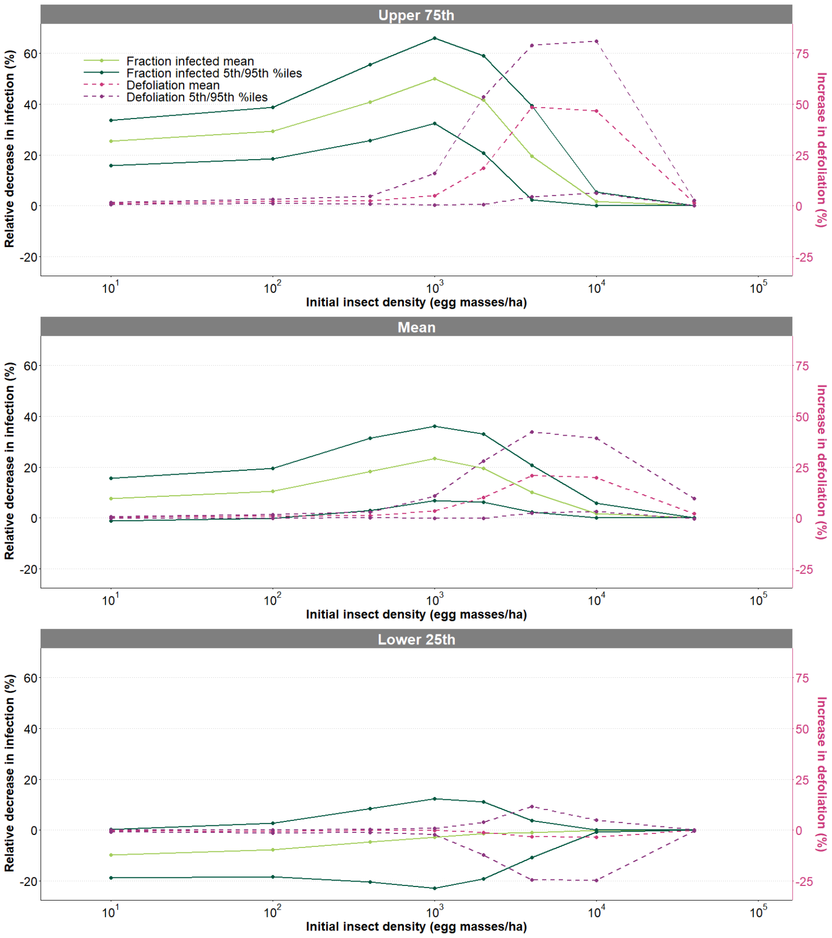

**Fig. 21** Summary plots of the effects of stochasticity on our model's projections. The middle plot is a modification of fig. 3 in the main text. Because we show the relative decrease in the fraction infected on the left vertical axis, the upper 75th and lower 25th percentiles indicate more severe and less severe effects of climate change respectively. Because changes at the 25th percentiles are only moderately less severe than the changes at the means, whereas changes at the 75th percentiles are substantially more severe than the changes at the means, a consideration of stochasticity leads to more pessimistic conclusions than the conclusions in the main text.

[4] Hajek, A. E. *et al.* Allozyme and restriction fragment length polymorphism analyses confirm entomophaga maimaiga responsible for 1989 epizootics in north american gypsy moth populations. *Proceedings of the National Academy of Sciences* **87**, 6979–6982 (1990).

[5] Hajek, A. E. *et al.* Host range of the gypsy moth (lepidoptera: Lymantriidae) pathogen entomophaga maimaiga (zygomycetes: Entomophthorales) in the field versus laboratory. *Environmental Entomology* **25**, 709–721 (1996).

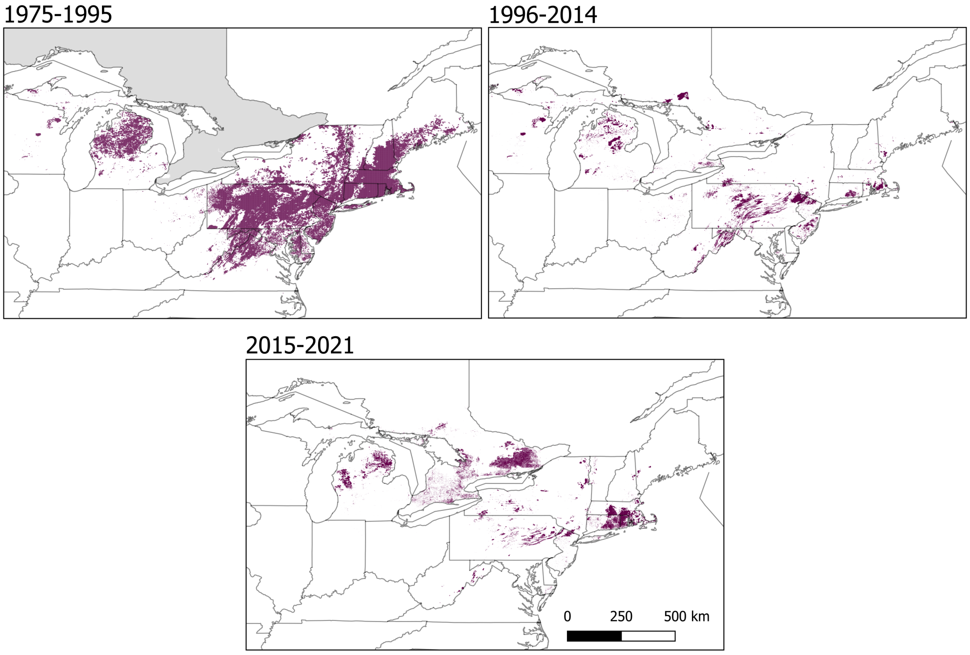

**Fig. 22** Comparison of defoliation maps for different periods. The gray shading indicates that data for Ontario are unavailable before 1996.

[11] Han, X. & Kloeden, P. E. *Random ordinary differential equations and their numerical solution* (Springer, 2017).

[28] Liebhold, A., Gottschalk, K., Luzader, E., Mason, D. & Bush, R. Gypsy moth in the united states: An atlas. forest service general technical report ne-233. Tech. Rep., Forest Service, Delaware, OH (United States). Forestry Sciences Lab. (1997).

[29] Régnière, J., Nealis, V. & Porter, K. in *Climate suitability and management of the gypsy moth invasion into canada* 135–148 (Springer, 2008).

[30] Burnham, K. P. & Anderson, D. R. *Model Selection and Multimodel Inference: A Practical Information-Theoretic Approach* (Springer, 2010).

[31] Thompson, L. M. *et al.* Climate-related geographical variation in performance traits across the invasion front of a widespread non-native insect. *Journal of biogeography* **48**, 405–414 (2021).

[32] Hajek, A. E., Diss-Torrance, A. L., Siegert, N. W. & Liebhold, A. M. Inoculative releases and natural spread of the fungal pathogen *entomophaga maimaiga* (entomophthorales: Entomophthoraceae) into us populations of gypsy moth, lymantria dispar (lepidoptera: Erebidae). *Environmental Entomology* **50**, 1007–1015 (2021).

[33] Elkinton, J. S. *et al.* Relating aerial deposition of entomophaga maimaiga conidia (zoopagomycota: Entomophthorales) to mortality of gypsy moth (lepidoptera: Erebidae) larvae and nearby defoliation. *Environmental Entomology* **48**, 1214–1222 (2019).
